# Deanticoagulated heparins drive a novel repair-secretory epithelial program to restore mucosal barrier in ulcerative colitis

**DOI:** 10.64898/2026.02.12.705304

**Authors:** Wei Hu, Zhen Kang Liu, Long Huang, Wen Zeng, Xiao Yi Ren, Yang Ji, Zheng Yang Song, Qin Xuan Zhou, Hai Hong Chen, Bing Xu, Can Yang Zhang, Chong Zhang, Peter E. Lobie, Zhen Qing Zhang, Yun Sheng Yang, Hong Chao Zhang, Ye Chen, Xuan Jiang, Yi Wang, Xin-Hui Xing

## Abstract

Epithelial mucus barrier dysfunction is a pathological hallmark of ulcerative colitis (UC), yet current clinical therapeutic strategies primarily suppress inflammation, lacking reliable approaches for restoration of the mucosal layer. Here, through a systematic screen of our established library of deanticoagulated heparins, we identified NALHP, a non-anticoagulant low-molecular-weight heparin derivative, and its representative fraction S6 as orally active epithelial repair-promoting glycans. NALHP/S6 restored crypt architecture, mucus production and epithelial barrier integrity in experimental colitis. In UC patient-derived colonic organoids, NALHP/S6 reduced aberrant stem/proliferative programs while promoting secretory, absorptive and junctional maturation. Single-cell transcriptomics and temporal validation identified a transient repair-secretory transitional state (RSTS) positioned between classical Wnt-associated stemness and mature barrier-forming epithelial states. Transcriptomic analysis revealed coordinated attenuation of Wnt and Notch signaling, whereas pharmacological reactivation of these pathways opposed NALHP/S6-induced epithelial state progression. These findings define mucosal repair as a regulated epithelial-state transition and identify NALHP/S6 as glycan-based modulators capable of restoring this progression in UC epithelium.

## Introduction

Ulcerative colitis (UC) is a chronic inflammatory disease characterized by recurrent mucosal injury and disruption of the colonic epithelial barrier^1–3^. Although suppression of inflammation remains central to current therapeutic strategies, durable disease control increasingly depends on achieving mucosal healing and restoration of epithelial function^4,5^. The colonic barrier is maintained not only by mucus-producing secretory cells but also by coordinated renewal of crypt progenitors, maturation of absorptive epithelial cells and formation of functional intercellular junctions^6^. Failure to restore this cellular organization may therefore persist even when inflammatory activity is reduced^7^.

Intestinal epithelial repair is a dynamic process involving transient changes in cellular identity rather than a simple replacement of lost terminally differentiated cells. Injury-responsive, regenerative and fetal-like epithelial programs have been described in damaged intestine, indicating that epithelial cells can enter intermediate states before re-establishing mature tissue architecture^8–10^. However, how such repair-associated states are organized in UC epithelium, how they relate to subsequent absorptive and secretory maturation, and whether defective progression through these states can be pharmacologically restored remain poorly understood.

Heparin (HP) is a highly sulfated glycosaminoglycan whose extended and structurally heterogeneous glycan backbone supports biological activities beyond anticoagulation^12–14^. Its canonical anticoagulant function depends on a defined antithrombin-binding pentasaccharide sequence, whereas the remaining non-anticoagulant domains can interact with diverse extracellular proteins and signaling systems. Previous attempts to use anticoagulant HP for UC treatment have produced inconsistent outcomes^12–14^ and are limited by bleeding risk and an incompletely defined structure–function relationship^15,16^. We therefore reasoned that a controlled removal of anticoagulant activity combined with activity-guided depolymerization of HP might uncover heparin-derived glycans capable of modulating epithelial repair independently of classical anticoagulation.

Here, we screened a library^17^ of deanticoagulated heparins and identified NALHP and its representative low-molecular-weight fraction S6 as orally active barrier-restorative glycans. Integrating experimental colitis, UC patient-derived organoids, single-cell transcriptomics and temporal validation, we find that NALHP/S6 do not simply expand a single differentiated lineage. Instead, its treatment promotes exit from an aberrant stem/proliferative program, transient engagement of a repair-secretory transitional state (RSTS), and subsequent progression toward complementary absorptive and mucus-barrier epithelial states. Transcriptomic profiling and bidirectional pathway perturbation further revealed that Wnt and Notch act as stage-specific regulatory gates of NALHP-induced repair, together defining the signaling logic that enables epithelial cells to progress through RSTS toward mature barrier-forming states.

## Results

### Activity-guided screening identifies NALHP and S6 as bioactive deanticoagulated heparin *derivatives*

We previously established a library of various structurally modified deanticoagulated HPs, including NAHP (non-anticoagulant HP), I –45 (prepared by enzymatic hydrolysis with heparinase I until the absorbance increased to 45), NAI45 (non-anticoagulant I –45), III, II –2-Odes-1 (prepared by combined enzymatic hydrolysis with heparinase III and heparinase II) ^17^, together with Eno (enoxaparin) and NAEno (non-anticoagulant enoxaparin) (Fig 1a). Screening in DSS-induced experimental colitis revealed heterogeneous therapeutic activity among these derivatives (Supplementary Fig. 1). Periodate and borohydride –mediated deanticoagulation generated NAHP with markedly reduced anti-Xa activity while retaining partial therapeutic activity (Fig. 1a, c and Supplementary Fig. 1). Controlled heparinase I depolymerization further generated NALHP and NAULHP, both of which showed stronger therapeutic phenotypes than parental NAHP in the DSS model (Supplementary Fig. 2). NALHP showed greater survival benefit and a more reproducible therapeutic phenotype than NAULHP and was therefore selected for subsequent investigation (Supplementary Fig. 2i and Fig. 1d–f).

**Fig 1.**
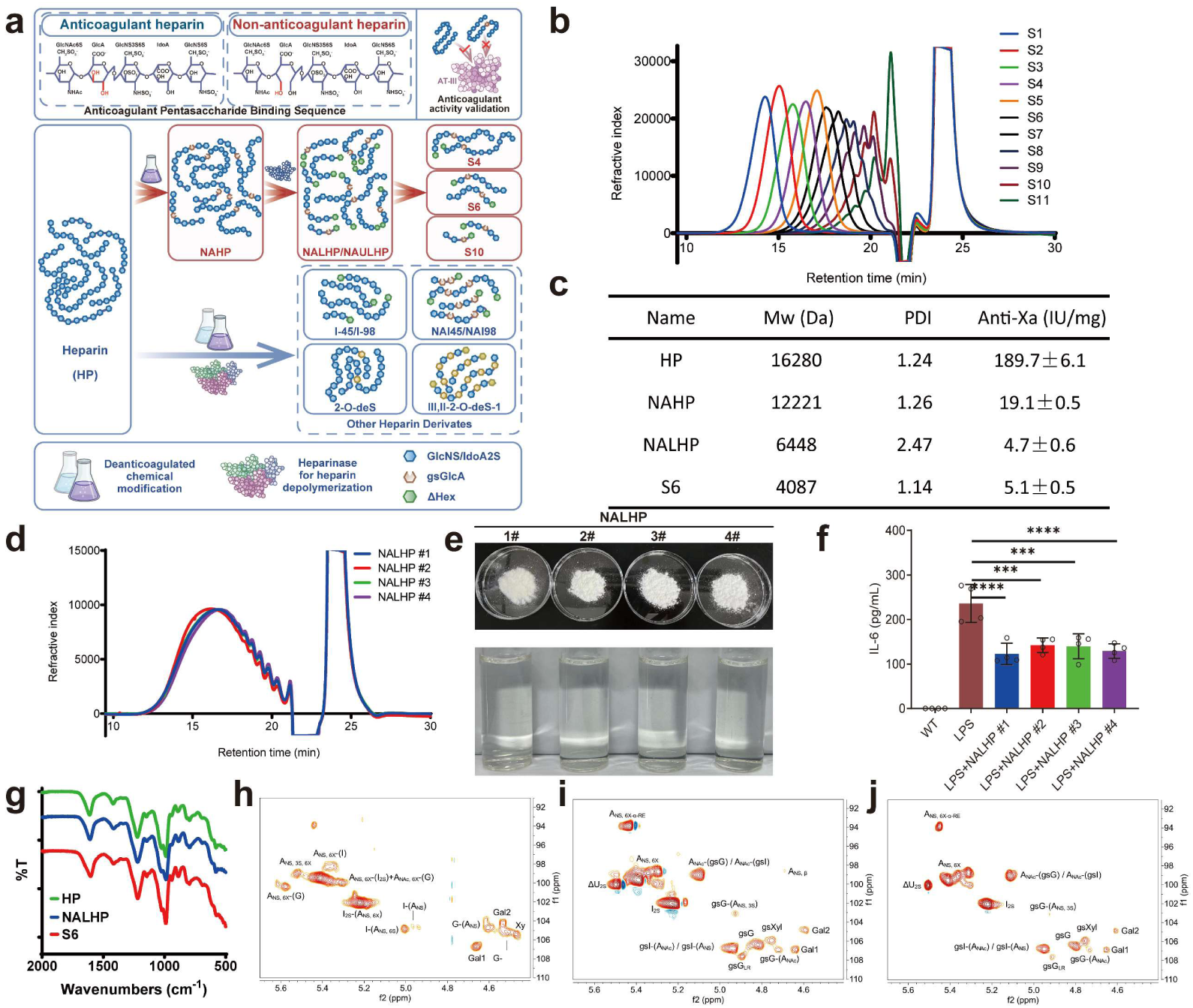
Physicochemical characterization identifies reproducible deanticoagulated HPs, NALHP and S6. **a**, A library of deanticoagulated HPs was created: NAHP (non-anticoagulant HP, prepared by periodate oxidation followed by sodium borohydride reduction of HP), I –45 (prepared by enzymatic hydrolysis with heparinase I until the absorbance increased to 45), NAI45 (non-anticoagulant I –45, prepared by periodate oxidation followed by sodium borohydride reduction of I –45), III, II –2-Odes-1 (prepared by combined enzymatic hydrolysis with heparinase III and heparinase II) ^17^, Eno (enoxaparin), and NAEno (non-anticoagulant Eno, prepared from enoxaparin). **b,** Size-exclusion chromatographic profiles of fractions S1–S11. **c,** Molecular weight (Mw), polydispersity index (PDI) and anti-factor Xa (anti-Xa) activity of native heparin (HP and deanticoagulated HPs. **d,** Size-exclusion chromatographic profiles of four independently prepared batches of NALHP, showing batch-to-batch consistency in molecular-weight distribution. **e,** Morphology and solubility of NALHP from the four production batches (#1–#4). **f,** Effects of independently prepared NALHP batches on LPS-induced IL-6 production in RAW264.7 cells, used as a biological activity quality-control assay. **g**, Representative Fourier-transform infrared spectra of HP, NALHP and S6. **h–j,** HSQC spectra of HP **(h)**, NALHP **(i)** and S6 **(j)**. A represents GlcN, G represents GlcA, I represents IdoA, ΔU represents ΔHexA; in the group with sulfonic acid modification, X indicates that the sulfonic acid group modification at this position is optional; the parentheses indicate that the polysaccharide is connected to the target sugar unit, not the chemically attributed target sugar unit; RE represents the reducing end; and LR represents the GlcA-Gal-Gal-Xyl tetrasaccharide structure of HP, which is where diverse protein serine residues attach to HP. The data are presented as the mean ± SD values; * *p* < 0.05, ** *p* < 0.01, *** *p* < 0.001, **** *p* < 0.0001, and ns (not significant) *p* > 0.05. Body-weight changes were analyzed using two-way repeated-measures ANOVA or a mixed-effects model, followed by the specified multiple-comparison test. Endpoint measurements were analyzed using ordinary one-way ANOVA followed by Dunnett’s multiple-comparison test.

Size-exclusion fractionation of NALHP generated 11 fractions spanning a broad molecular-weight range (S1–S11) (Fig. 1b and Supplementary Table 1). Integrated evaluation of body weight, colon length, spleen index, inflammatory cytokines and histopathological injury showed enrichment of therapeutic activity within intermediate low-molecular-weight fractions (Supplementary Fig. 3). S6 had an average molecular weight of approximately 4.1 kDa, a narrow molecular-weight distribution and minimal anti-Xa activity (Fig. 1c and Supplementary Table 1). S6 reproduced the major therapeutic phenotype of parental NALHP and was therefore selected as a representative bioactive NALHP fraction (Supplementary Fig. 3). These activity-guided analyses identified NALHP and S6 as reproducible, deanticoagulated heparin-derived preparations with prominent therapeutic activity in experimental colitis (Fig. 1 and Supplementary Figs. 1-3).

### NALHP and S6 retain reproducible heparin-like structural features after deanticoagulation and depolymerization

Independent production batches of NALHP showed closely overlapping size-exclusion chromatographic profiles and comparable physicochemical properties (Fig. 1d, e and Supplementary Tables 3–5). The independently prepared batches also showed comparable effects in the RAW264.7 biological quality-control assay (Fig. 1f). FTIR spectroscopy showed that NALHP and S6 retained the major sulfate– and glycosidic-bond-associated features of the parental heparin structure after chemical deanticoagulation and controlled depolymerization (Fig. 1g, Supplementary Fig. 4 and Supplementary Table 2). ^-1^H NMR and ^-13^C NMR provided additional evidence for conservation of the major heparin-derived structural framework (Supplementary Figs. 5 and 6 and Supplementary Tables 6 and 7). HSQC spectra further showed broadly related structural signatures among NALHP and the bioactive S4 and S6 fractions (Fig. 1h–j and Supplementary Fig. 7).

In contrast, the lower-molecular-weight S10 fraction showed loss or alteration of several linkage-region and reducing-end-associated signals relative to NALHP/S6 (Supplementary Fig. 7). Independent preparative separations reproduced the chromatographic isolation of S4, S6 and S10, supporting the robustness of the fractionation procedure (Supplementary Fig. 8). Thus, therapeutic activity is associated with a reproducible molecular-weight and structural context rather than with nonspecific depolymerization of the heparin backbone (Fig. 1 and Supplementary Figs. 4-8).

### Oral NALHP and S6 restore epithelial barrier architecture in experimental colitis

Oral NALHP and S6 attenuated DSS-associated body-weight loss and restored colon length relative to untreated DSS mice (Fig. 2b, c and e). Both treatments also reduced the DSS-induced increase in spleen index (Fig. 2d, f). Serum IL-6 and TNF-α concentrations decreased following NALHP or S6 treatment, demonstrating concomitant improvement of the inflammatory environment (Fig. 2g, h).

**Fig 2.**
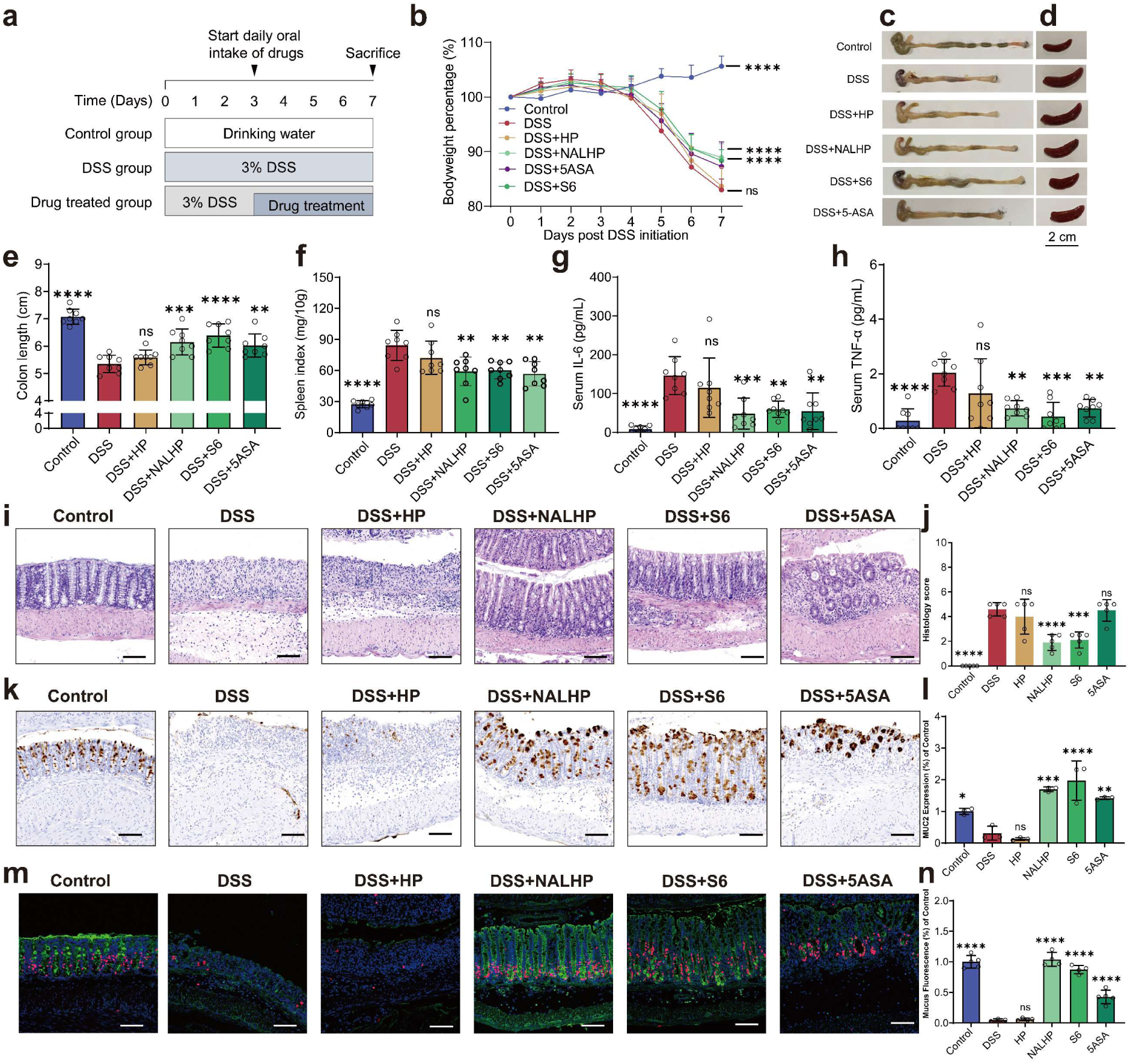
NALHP and S6 restore epithelial barrier architecture in experimental colitis. **a**, DSS (3%) was added to the drinking water for 7 days to induce acute colitis in mice (n = 8). The mice in the control group were provided regular drinking water (n = 8), and the mice in the drug treatment group were administered 30 mg/kg HP, NALHP, S6, or 5-ASA on the third day (n = 8). **b,** Changes in body weight. **c–f,** Mice were sacrificed on day 7, and the colon (**c**) and spleen (**d**) were harvested. The colon length (**e**) and the spleen index (**f**) were quantified. **g–h,** The serum concentrations of IL-6 (**g**) and TNF-α (**h**) were measured by ELISA after the mice were sacrificed on day 7. **i-j,** Representative images of colons stained with hematoxylin and eosin(**i)** and (**j)** the quantitative histological scores; scale bar, 100 μm. **k-l**, Representative images of colons stained with hematoxylin and MUC2(**k)** and (**l)** the quantitative immunohistochemical scores; scale bar, 100 μm. **m-n,** Representative images of colons stained with EdU (red) and WGA (green) (**m)** and (**n)** the quantitative immunohistochemical scores of mucus barrier/EdU quantification; scale bar, 100 μm. Representative images or quantitative analyses of n = 8 animals (**b–h**) and n = 5 biologically independent samples (**i–n**) are shown. The data are presented as the mean ± SD values; * *p* < 0.05, ** *p* < 0.01, *** *p* < 0.001, **** *p* < 0.0001, and ns (not significant) *p* > 0.05. The data were analyzed by an ordinary one-way ANOVA with multiple comparison testing.

DSS exposure caused marked crypt disruption and epithelial injury, whereas NALHP and S6 restored crypt organization and reduced histopathological injury scores (Fig. 2i, j). MUC2-positive epithelial staining was substantially increased after NALHP/S6 treatment (Fig. 2k, l). WGA staining further demonstrated restoration of mucus-associated glycoprotein coverage (Fig. 2m, n). EdU staining showed recovery of proliferative activity within regenerating colonic epithelium (Fig. 2m, n).

Native anticoagulant HP did not reproduce the same epithelial-repair phenotype, whereas 5-ASA produced only partial restoration under the experimental conditions used here (Fig. 2i–n). Repeated oral administration of NALHP to healthy mice produced no overt changes in the measured body-weight, gross-organ or colonic histological parameters (Supplementary Fig. 9). Alternative systemic administration routes did not improve S6 efficacy relative to oral administration (Supplementary Fig. 10). Together, these data show that orally administered NALHP and S6 restore the structural and functional components of the colonic epithelial barrier in DSS-induced colitis while simultaneously reducing inflammatory activity (Fig. 2 and Supplementary Figs. 3 and 10).

### Epithelial repair more closely tracks therapeutic efficacy than systemic immune or microbiome remodeling

Several deanticoagulated heparin derivatives altered intestinal microbial composition toward profiles observed in control mice despite showing different degrees of therapeutic efficacy (Supplementary Fig. 11a–c). Metabolomic remodeling was similarly observed across multiple derivatives and was not uniquely associated with the NALHP/S6 therapeutic phenotype (Supplementary Fig. 11d–f). Thus, microbiome and metabolome changes were not sufficient to explain the differential efficacy of NALHP and S6 (Supplementary Fig. 11).

In contrast, colonic transcriptomic responses associated with the more active NALHP-derived preparations were enriched for epithelial proliferation, wound repair, cell-fate regulation, cell-junction organization and Wnt-associated biological processes (Supplementary Fig. 12). NALHP and S6 also reduced colonic CD11b-associated inflammatory infiltration and P-STAT3 staining (Supplementary Fig. 13). Splenic CD3-and CD4-associated staining patterns showed comparatively limited treatment-associated alterations (Supplementary Fig. 14). NALHP/S6 treatment reorganized PCNA-positive proliferative cells within the regenerating crypts and altered TCF1/7-associated signaling (Supplementary Fig. 15). P-ERK localization was also remodeled in treated crypts (Supplementary Fig. 16). Across the tested derivatives, therapeutic efficacy showed a closer association with host epithelial repair programs than with a uniform systemic immune, microbiome or metabolome response.

### NALHP and S6 reproduce broad epithelial maturation in UC patient-derived organoids

UC patient-derived colon organoids recapitulated the goblet cell deficiency observed in diseased colonic mucosa. (Supplementary Fig. 17a, b). Therefore, we established a UC organoid model based on colon epithelial cells isolated from the colon mucosal layer of Chinese patients diagnosed with UC (Supplementary Fig. 18a). NALHP and S6 induced reproducible morphological remodeling of UC patient-derived colonic organoids, whereas 5-ASA produced little comparable morphological change (Fig. 3a, b). Conventional single-cell annotation showed that untreated UC organoids were dominated by stem and transit-amplifying populations (Fig. 3c, d and Supplementary Fig. 18). NALHP and S6 markedly reduced these proliferative populations while increasing multiple differentiated epithelial populations.

**Fig 3.**
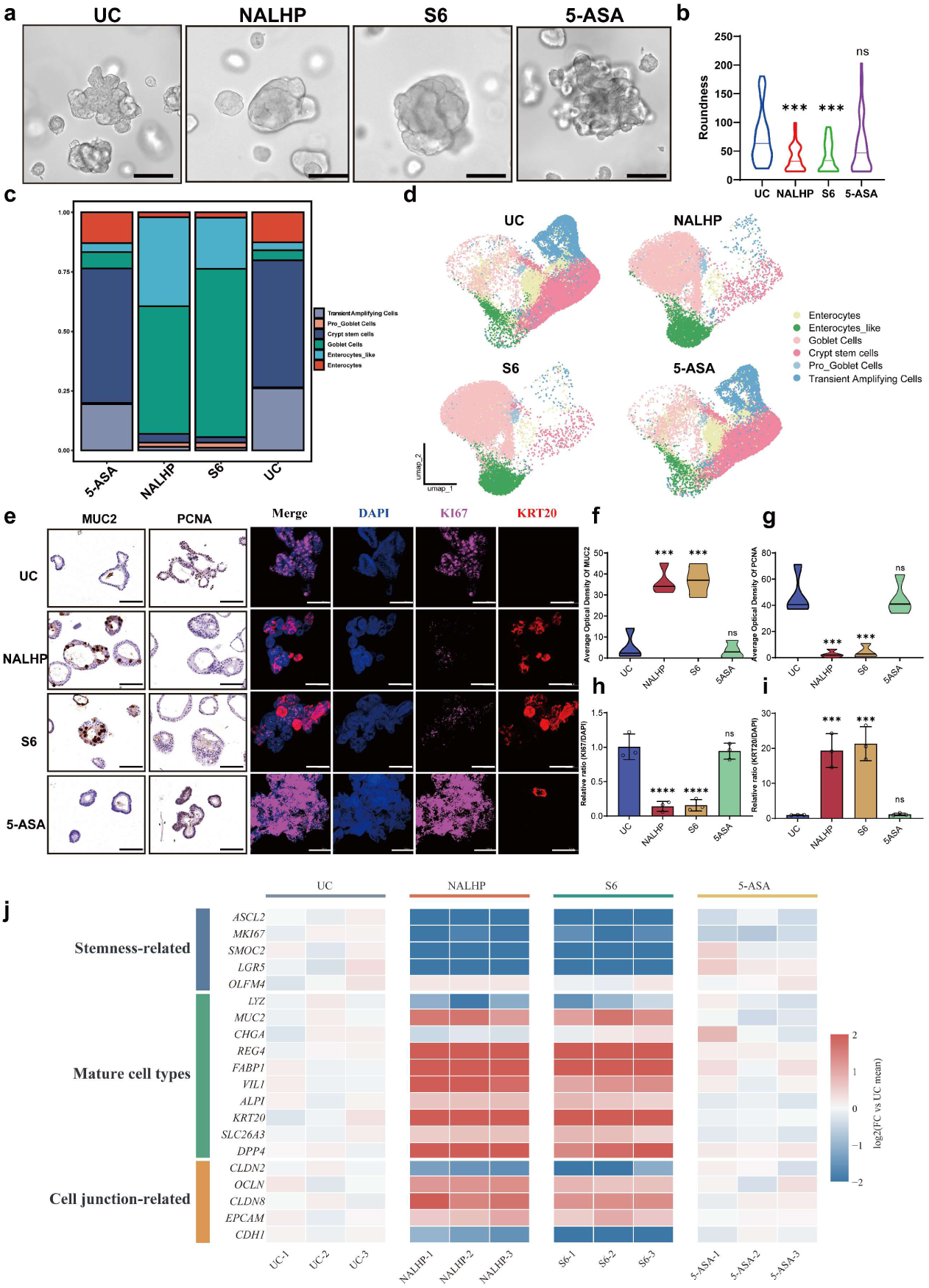
NALHP and S6 reproduce epithelial repair and functional maturation in UC patient-derived organoids. **a**, Representative bright-field images of untreated UC patient-derived colonic organoids and organoids treated with NALHP, S6 or 5-ASA. Scale bars, 200 μm. **b**, Quantification of organoid roundness after treatment. **c**, Relative proportions of conventional epithelial cell types identified by single-cell RNA sequencing, including crypt stem cells, transit-amplifying cells, pro-goblet cells, goblet cells, enterocyte-like cells and mature enterocytes. **d,** UMAP visualization of epithelial cells from UC, NALHP-, S6– and 5-ASA-treated organoids, colored by conventional epithelial cell identity. **e**, Representative MUC2 and PCNA immunohistochemical staining and KI67/KRT20 immunofluorescence in UC organoids following the indicated treatments. Scale bars, 50 μm. **f-g**, Quantification of MUC2 (**f**) and PCNA (**g**) staining. **h-i**, Quantification of KI67-positive (**h**) and KRT20-positive (**i**) epithelial cells relative to DAPI-positive nuclei. **j**, Heatmap showing changes in stemness-associated, functionally mature epithelial and cell-junction-associated genes across three independently derived UC organoid lines after NALHP, S6 or 5-ASA treatment. Expression values are shown as log2 fold change relative to the mean of the untreated UC group. Data are presented as mean ± SD where applicable; n = 3 biologically independent organoid cultures. Statistical significance was determined using an ordinary one-way ANOVA; * *p* < 0.05, ** *p* < 0.01, *** *p* < 0.001, **** *p* < 0.0001 and ns (not significant) *p* > 0. 05.

MUC2 staining increased substantially after NALHP/S6 treatment, demonstrating enhanced mucus-associated maturation (Fig. 3e, f). PCNA and KI67-associated proliferative features decreased after treatment (Fig. 3e, g and h). KRT20-positive mature epithelial cells increased concurrently (Fig. 3e, i).

Transcript-level analysis across three independently derived UC organoid lines showed broad downregulation of stemness-associated genes and induction of mature epithelial programs after NALHP/S6 treatment (Fig. 3j). The induced mature programs included mucus-associated genes as well as absorptive markers such as *KRT20, SLC26A3, FABP*-family genes, *VIL1, ALPI* and *DPP4* (Fig. 3j). Junction-associated genes, including epithelial adhesion and tight-junction-related transcripts, were also increased following NALHP/S6 treatment (Fig. 3j).

Patient-level single-cell maps demonstrated comparable epithelial remodeling across three independent UC organoid lines (Supplementary Fig. 19). NALHP-associated restoration of junctional ZO-1 organization was independently observed in cytokine-challenged colonic epithelial cells and UC organoids (Supplementary Figs. 20 and 21). These findings demonstrate that NALHP/S6 do not selectively induce a single terminal lineage but instead suppress an abnormal stem/proliferative program while promoting secretory, absorptive and junctional epithelial maturation.

### A transient repair-secretory state precedes epithelial barrier maturation

To explore the pronounced differentiation phenotype induced by NALHP/S6, we first performed pseudotime analysis using the conventional epithelial cell annotations. The trajectory indicated that the differentiated epithelial populations observed after NALHP/S6 treatment originated from the crypt stem/transit-amplifying compartment (Fig. 4a, b and Supplementary Fig. 22). We next examined the differentiation process in real time using Day1, Day3 and Day5 qPCR. Whereas stemness-associated genes generally declined and mature epithelial markers increased during treatment, a subset of secretory-associated genes, such as *SPDEF* and *MUC2*, showed a distinct transient pattern, increasing during the early-to-intermediate phase and subsequently decreasing (Fig. 4c).

**Fig 4.**
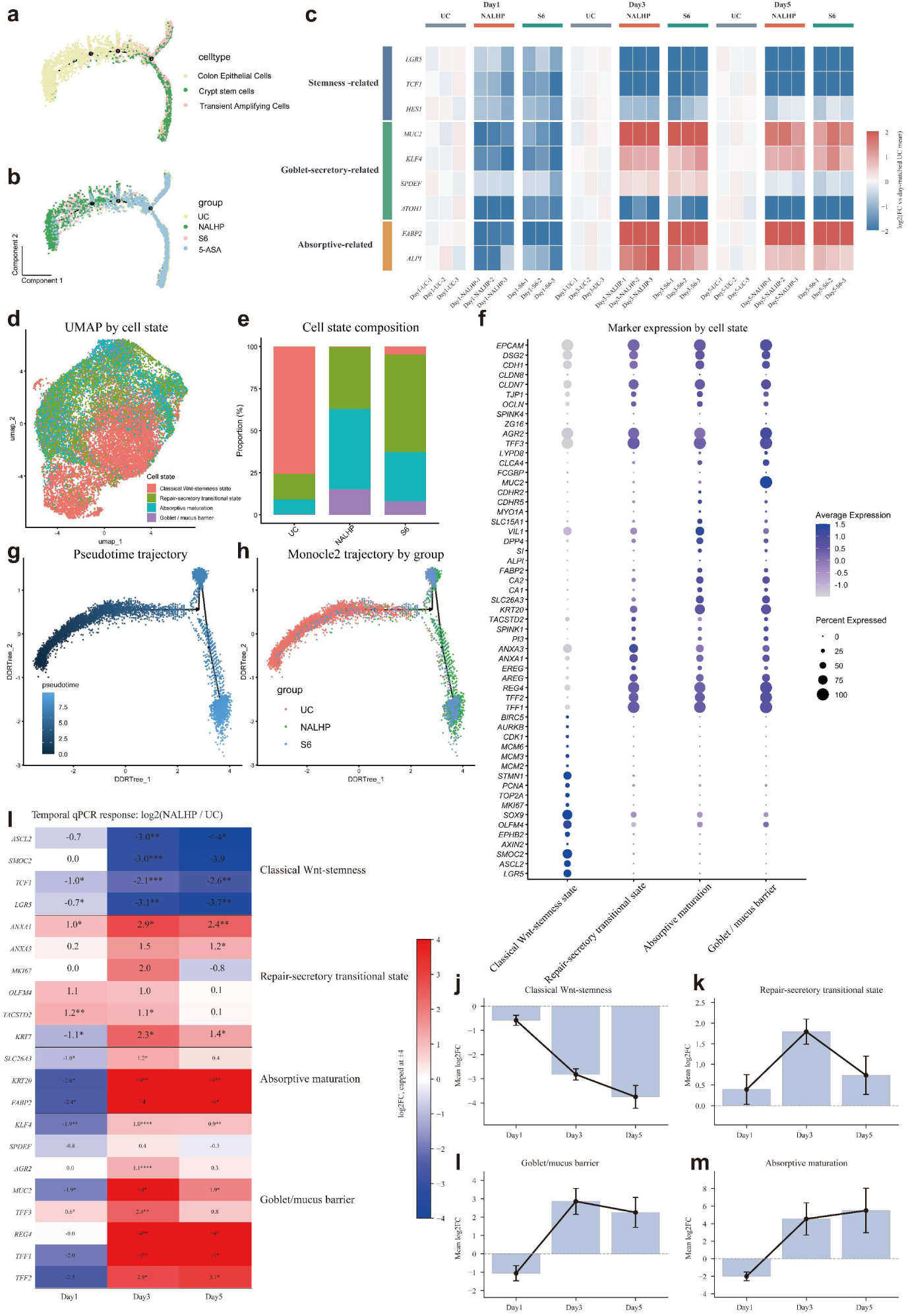
A transient repair-secretory state links abnormal stemness to epithelial barrier maturation. **a-b**, Monocle 2 pseudotime projection of epithelial cells colored by cell type (**a**) and treatment group (**b**). **c,** Temporal expression profiles of representative stemness-associated, goblet-secretory and absorptive maturation genes in UC organoids treated with NALHP or S6 at days 1, 3 and 5. Expression values are shown relative to day-matched untreated UC organoids. **d**, UMAP visualization of four epithelial states defined from single-cell transcriptomes: classical Wnt-stemness, repair-secretory transitional state (RSTS), absorptive maturation and goblet/mucus-barrier state. **e**, Relative abundance of the four epithelial states in untreated UC, NALHP– and S6-treated organoids. **f**, Dot plot showing representative marker genes defining each epithelial state; dot size represents the percentage of cells expressing each gene and color denotes average expression. **g**, Monocle 2 pseudotime reconstruction of epithelial-state progression. **h**, The same trajectory colored by treatment group, showing redistribution of NALHP– and S6-treated cells along the differentiation trajectory. **i**, Temporal qPCR heatmap showing log2 fold changes in representative genes after NALHP treatment relative to day-matched UC controls. Genes are grouped according to classical Wnt-stemness, RSTS, absorptive maturation and goblet/mucus-barrier programs. **j–m**, Mean temporal changes in the classical Wnt-stemness (**j**), RSTS (**k**), goblet/mucus-barrier (**l**) and absorptive maturation (**m**) gene modules. RSTS exhibited a transient intermediate response, whereas downstream absorptive and mucus-barrier programs remained elevated during later stages of treatment. Representative results of the quantitative analysis of n = 3 biologically independent samples are shown (**c, l and n**). The data are presented as the mean ± SD values; * *p* < 0.05, ** *p* < 0.01, *** *p* < 0.001, **** *p* < 0.0001, and ns (not significant) *p* > 0.05. The data were analyzed by an ordinary one-way ANOVA.

This temporal pattern suggested that NALHP/S6-induced epithelial maturation might involve an intermediate repair-associated program rather than a direct transition from stem/proliferative cells to mature epithelial cells. To investigate this intermediate state, we re-analyzed the single-cell transcriptomic data in the context of intestinal epithelial injury and repair. Four transcriptional states were resolved: classical Wnt-stemness, a repair-secretory transitional state (RSTS), absorptive maturation and a goblet/mucus-barrier state (Fig. 4d–f). RSTS was characterized by repair– and secretory-associated genes, including *TFF1, TFF2, REG4, AREG, EREG, ANXA1, ANXA3, PI3, SPINK1, TACSTD2* and *KRT7*, whereas the two downstream mature states were characterized by absorptive or mucus-barrier-associated programs, respectively (Fig. 4f). NALHP and S6 reduced the proportion of cells in the classical Wnt-stemness state and redistributed epithelial cells toward RSTS and the downstream mature states (Fig. 4d, e).

Reconstruction of the trajectory using the refined state annotation positioned RSTS between classical Wnt-stemness and the mature absorptive and goblet/mucus-barrier states (Fig. 4g, h). We subsequently reorganized the independent Day1–Day5 temporal qPCR dataset according to these four state programs. Classical Wnt-stemness progressively declined, whereas the RSTS module increased to a maximum around Day3 and subsequently decreased; in contrast, absorptive and goblet/mucus-barrier programs increased and remained elevated during the later phase of treatment (Fig. 4i–m and Supplementary Fig. 23).

Together, the initial pseudotime analysis, real-time observation of a transient intermediate transcriptional response, and subsequent state-resolved single-cell re-analysis support a model in which NALHP/S6 drive UC epithelium from abnormal stemness through a transient repair-secretory state toward mature barrier-forming epithelial states (Fig. 4 and Supplementary Figs. 22 and 23).

### Transcriptomic remodeling links epithelial state progression to Wnt and Notch attenuation

Differential-expression analysis showed broad suppression of stemness-associated genes and induction of repair– and maturation-associated genes in NALHP-treated UC organoids (Fig. 5a). Gene-set enrichment analysis identified negative enrichment of Wnt– and Notch-associated signaling and positive enrichment of tight-junction-related programs following NALHP treatment (Fig. 5b and Supplementary Fig. 24a–f). MAPK and JAK–STAT signaling did not show the same enrichment pattern under these conditions (Fig. 5b and Supplementary Fig. 24a–f).

**Fig 5.**
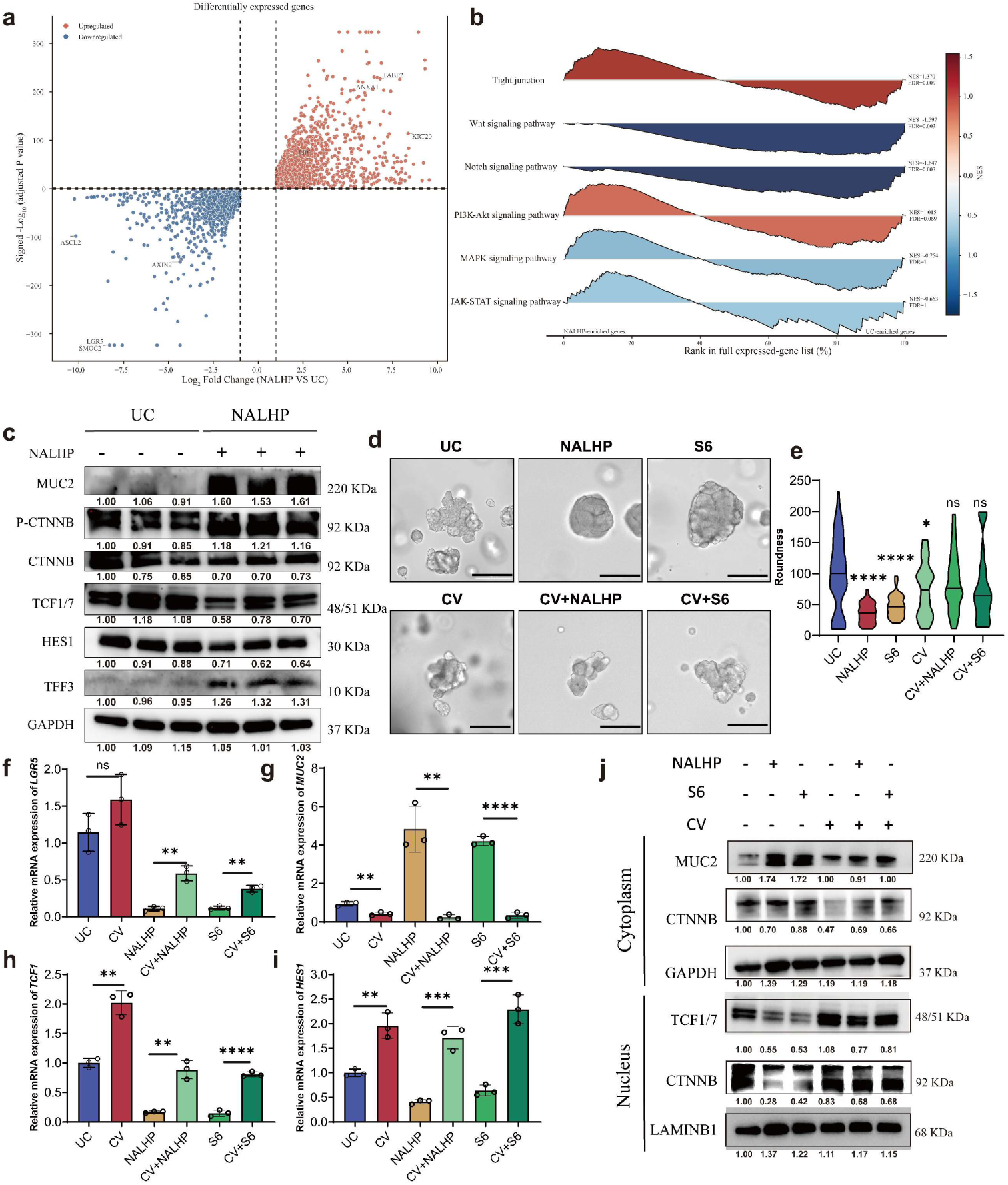
Wnt and Notch reactivation resists NALHP-driven epithelial state progression. **a**, Volcano plot of differentially expressed genes between NALHP-treated and untreated UC organoids. Representative genes associated with classical stemness, epithelial repair and maturation are indicated. **b**, Gene-set enrichment analysis of epithelial and signaling pathways in NALHP-treated versus untreated UC organoids. Positive normalized enrichment scores (NES) indicate enrichment in NALHP-treated organoids and negative NES values indicate enrichment in untreated UC organoids. Tight-junction-associated genes were positively enriched, whereas Wnt and Notch signaling pathways were negatively enriched following NALHP treatment. **c**, Immunoblot analysis of MUC2, phosphorylated β-catenin (p-CTNNB), total β-catenin (CTNNB), TCF1/7, HES1 and TFF3 in untreated and NALHP-treated UC organoids. GAPDH served as a loading control. **d**, Representative bright-field images of UC organoids treated with NALHP or S6 in the absence or presence of combined Wnt and Notch reactivation (CV). CV denotes combined treatment with the Wnt activator Laduviglusib (CHIR99021, C) and the Notch activator sodium valproate (VPA, V). Scale bars, 200 μm. **e**, Quantification of organoid roundness under the indicated conditions. **f–i**, Relative mRNA expression of LGR5 (**f**), MUC2 (**g**), TCF1 (**h**) and HES1 (**i**) following NALHP/S6 treatment with or without Wnt/Notch reactivation. **j**, Cytoplasmic and nuclear immunoblot analysis of MUC2, CTNNB and TCF1/7 following the indicated treatments. Numbers below individual bands indicate relative densitometric intensities normalized to GAPDH for cytoplasmic proteins or LAMINB1 for nuclear proteins, with the corresponding UC value set to 1.0. Combined Wnt/Notch reactivation partially restored stemness-associated signaling and opposed NALHP/S6-induced epithelial maturation. Data are presented as mean ±SD; n =3 biologically independent experiments. The data are presented as the mean ± SD values; * *p* < 0.05, ** *p* < 0.01, *** *p* < 0.001, **** *p* < 0.0001, and ns (not significant) *p* > 0.05. The data were analyzed by an ordinary one-way ANOVA.

Immunoblot analysis showed corresponding changes in β-catenin/TCF-associated Wnt output, HES1 and mature epithelial proteins following NALHP treatment (Fig. 5c). These unbiased transcriptomic and protein-level analyses identified coordinated changes in Wnt– and Notch-associated signaling as candidate regulatory features of NALHP-driven epithelial state progression (Fig. 5a–c and Supplementary Fig. 24a–f).

### Wnt and Notch reactivation functionally constrains distinct steps of NALHP-driven epithelial state progression

Combined Wnt and Notch inhibition with IWP-2 and DAPT altered organoid morphology, reduced LGR5 expression and increased MUC2 expression, thereby phenocopying selected features of the NALHP/S6 response (Supplementary Fig. 24g–i). Combined reactivation of Wnt and Notch signaling with CHIR99021 and valproic acid opposed NALHP/S6-induced morphological remodeling (Fig. 5d, e). Pathway reactivation partially restored LGR5, TCF1 and HES1 expression and reduced the induction of MUC2 (Fig. 5f–i). Nuclear and cytoplasmic immunoblot analysis further showed recovery of β-catenin/TCF-associated signaling under combined reactivation conditions (Fig. 5j).

Separate Wnt reactivation with CHIR99021 preferentially restored LGR5 and TCF1-associated outputs without comparably restoring Notch-associated HES1 signaling (Supplementary Fig. 25a– e). In contrast, Notch reactivation with valproic acid preferentially restored HES1 and opposed MUC2 induction without comparably restoring the Wnt-associated stemness program (Supplementary Fig. 25f–j).

These bidirectional perturbation experiments demonstrate that Wnt– and Notch-associated signaling functionally constrains NALHP/S6-driven epithelial state progression (Fig. 5d–j and Supplementary Figs. 23 and 25). The pathway-specific reactivation experiments further support complementary functions in which Wnt-associated signaling primarily constrains stemness exit, whereas Notch-associated signaling preferentially constrains subsequent secretory maturation (Supplementary Fig. 25). These data establish functional involvement of Wnt and Notch signaling but do not establish direct molecular binding of NALHP/S6 to a defined component of either pathway (Fig. 5 and Supplementary Fig. 25).

## Discussion

This study identifies a noncanonical function of heparin-derived glycans in epithelial tissue repair. Through chemical deanticoagulation, controlled enzymatic depolymerization, and activity-guided fractionation, we identified NALHP and its representative fraction S6 as orally active, non-anticoagulant heparin derivatives that restore the colonic mucus barrier in experimental colitis. Their activity was associated with improved crypt architecture, increased goblet cell abundance and MUC2 expression, and a shift in ulcerative colitis patient-derived organoids from stem/transit-amplifying states towards broad functional maturation encompassing absorptive, mucus-secretory and junctional programs. Single-cell trajectory reconstruction further positioned a repair-secretory transitional state (RSTS) between classical Wnt-associated stemness and mature barrier-forming epithelial states, while Day1–Day5 temporal measurements showed that this program rose transiently before declining as absorptive and mucus-barrier programs became established. Together, these observations support a model in which NALHP/S6 restore epithelial function by promoting progression through a temporally organized repair trajectory rather than by directly forcing crypt stem cells into a single terminal fate.

This interpretation is consistent with an emerging view that intestinal repair involves extensive epithelial plasticity and transient departures from homeostatic cell identities. YAP/TAZ-dependent reprogramming of damaged colonic epithelium can induce a regenerative state distinct from normal homeostasis, and injury can mobilize transient revival stem cells or colitis-associated regenerative stem-cell populations that contribute to tissue reconstruction^8,9,18^. Fetal-like reversion has likewise been observed after intestinal injury, demonstrating that regeneration can involve temporary acquisition of developmental programs rather than simple expansion of pre-existing LGR5-positive stem cells^19^. Human single-cell studies further show extensive epithelial remodeling in IBD, including altered stem-cell programs, depletion or redistribution of differentiated epithelial populations and emergence of inflammation-associated epithelial states^20–22^. Thus, RSTS should not currently be interpreted as an entirely independent or universally novel epithelial lineage. Instead, our data strongly support RSTS as a UC-associated repair configuration with prominent secretory and injury-response features that transiently occupies the trajectory between abnormal stemness and barrier maturation.

The temporal behavior of RSTS provides an important distinction from a stable regenerative population. RSTS-associated genes increased during the intermediate phase of NALHP treatment and declined thereafter, whereas mature absorptive and mucus-barrier modules remained elevated at later time points. This temporal organization was concordant with pseudotime reconstruction, which placed RSTS between classical Wnt-stemness and differentiated epithelial endpoints. Long-term colonic stem-cell culture studies have previously demonstrated cyclical transitions between homeostatic and injury-associated regenerative programs, supporting the concept that successful tissue repair requires not only entry into a regenerative state but also subsequent resolution and redifferentiation^8^. Our observations extend this principle to UC patient-derived epithelium by suggesting that NALHP does not simply activate repair-associated transcription but promotes progression beyond this intermediate state toward mature epithelial functions. Whether RSTS corresponds to a specific branch of previously described fetal-like, revival-stem-cell or metaplastic programs will require direct cross-dataset signature comparison and validation in active and healing human UC mucosa.

The organoid experiments also distinguish epithelial-state remodeling from a purely anti-inflammatory explanation for NALHP activity. NALHP/S6 reduced inflammatory cytokines and local P-STAT3– and CD11b-associated signals in experimental colitis, but major splenic T-cell-associated patterns were comparatively less altered. More importantly, reduction of the stem/proliferative program and induction of absorptive, secretory and junctional maturation were reproduced in patient-derived UC organoids, which lack circulating immune populations and an intact luminal microbiota. Human UC single-cell studies have shown that epithelial compartments themselves undergo substantial disease-associated transcriptional rewiring, supporting the biological relevance of examining epithelial-intrinsic programs independently of the full inflammatory environment^20,22^. In our experiments, 5-ASA also failed to reproduce the broad NALHP/S6-associated epithelial-state redistribution under the same organoid conditions. These findings do not imply that inflammation is dispensable for NALHP activity; rather, they indicate that suppression of inflammation alone is insufficient to account for the observed epithelial phenotype^23–25^.

Wnt and Notch signaling provide a plausible regulatory framework for this state progression^26–29^. LGR5 identifies rapidly cycling intestinal stem cells at the crypt base, and maintenance of this stem-cell compartment is closely coupled to high Wnt activity^30^. Notch signaling provides a complementary lineage-control mechanism: genetic or pharmacological inhibition of Notch converts proliferative crypt cells toward secretory, particularly goblet-cell, fates^31^, and cooperative interactions between Wnt and Notch regulate intestinal progenitor proliferation and differentiation^32^. Consistent with this framework, unbiased GSEA in our study identified coordinated negative enrichment of Wnt– and Notch-associated programs following NALHP treatment, whereas tight-junction-associated genes were positively enriched. Protein-level measurements showed corresponding changes in β-catenin/TCF-associated Wnt output and HES1, together with increased epithelial maturation proteins.

Importantly, functional perturbation moved the Wnt/Notch evidence beyond transcriptional association. Combined inhibition of Wnt and Notch signaling with IWP-2 and DAPT reproduced selected features of the NALHP phenotype, including reduced *LGR5* and increased *MUC2* expression. Conversely, combined reactivation with CHIR99021 and valproic acid opposed NALHP/S6-induced epithelial remodeling and partially restored *LGR5, TCF1 and HES1* while suppressing MUC2 induction. Separate pathway reactivation further indicated non-equivalent functions: Wnt reactivation preferentially restored *LGR5* and *TCF1*-associated outputs, whereas Notch reactivation preferentially restored *HES1* and opposed *MUC2* induction. These results support complementary regulatory roles in which Wnt-associated signaling primarily constrains exit from the stemness program, whereas Notch-associated signaling restricts subsequent secretory maturation. They do not, however, establish NALHP or S6 as direct inhibitors of a defined Wnt or Notch component; the immediate glycan-interacting proteins and upstream signaling events remain unresolved.

The physicochemical data further indicate that epithelial repair activity cannot be explained simply by removal of the canonical anticoagulant function of heparin. NALHP and S6 retained very low anti-Xa activity while preserving pronounced epithelial-repair activity, whereas the fractions with different molecular-weight and structural characteristics showed reduced activity. This interpretation is strengthened by the fondaparinux experiment: neither fondaparinux nor its deanticoagulated derivative reproduced the complete NALHP-associated pattern of MUC2, LGR5, PCNA and FABP1 expression in UC organoids (Supplementary Fig. 26). Thus, loss of anticoagulant function alone is insufficient to generate the epithelial phenotype. Rather, the activity is likely to depend on a combination of chain length, sulfation context, terminal structure and preservation of specific non-anticoagulant glycan domains. Because S6 remains a heterogeneous glycan fraction rather than a sequence-defined oligosaccharide, these data should be regarded as a structure–activity framework rather than identification of a single pharmacophore.

Several limitations define the current boundaries of this model. First, the transitional relationship between RSTS and mature epithelial states is supported by concordant pseudotime and true temporal expression data, but lineage tracing or directional single-cell approaches would provide stronger evidence of state ancestry. Second, the human evidence is currently derived predominantly from patient-derived organoids; direct demonstration of RSTS in active, repairing and healed UC mucosa will be important for establishing its in vivo human relevance. Third, although bidirectional pharmacological perturbation demonstrates functional involvement of Wnt and Notch signaling, the direct molecular target and earliest signaling event triggered by NALHP/S6 remain unknown. Finally, in vivo efficacy was established primarily in DSS-induced colitis, and additional chronic or immune-mediated models will be needed to determine whether restoration of this epithelial trajectory generalizes across distinct forms of intestinal injury. Nevertheless, the convergence of in vivo barrier restoration, human organoid remodeling, single-cell trajectory analysis, true temporal validation and pathway perturbation supports a model in which mucosal healing can be understood as a regulated sequence of epithelial-state transitions that can be pharmacologically restored by non-anticoagulant heparin-derived glycans.

## Methods and Materials

### Preparation of deanticoagulated HPs ^17,33,34^

The preparation process of deanticoagulated HPs was described previously. Briefly, pharmaceutical-grade HP sodium was purchased from Changshan Biochemical Pharmaceutical Co., Ltd. (Hebei, China). To eliminate anticoagulant activity, HP was dissolved in water at a concentration of 33 mg/ml. The solution was reacted with an equal volume of 0.2 M periodate sodium solution at 4 °C for 22 hr prior to neutralization reactions with 4 ml of glycol and 1.4 g of sodium borohydride per gram of HP at 4 °C for 1 hr and 16 hr, respectively. The resulting solution was adjusted to pH 7.0, filtered through a membrane, desalted using a 3 kDa Pellicon 2 Mini Cassette (Millipore, Germany), and lyophilized to obtain nonanticoagulated HP (NAHP). NAHP was subsequently subjected to further enzymolysis with recombinant heparinase I (Beijing Bio-Cell Technology Co., Ltd., China). NAHP was dissolved in a reaction buffer consisting of 20 mM Tris, 200 mM NaCl, and 5 mM CaCl_2_ at pH 7.4. The increase in the absorbance at 232 nm was recorded immediately after the heparinase I was added to the NAHP solution. Low-molecular-weight NAHP (NALHP) and ultra-low-molecular-weight NAHP (NAULHP) were obtained when the absorbance increased to 45 and 100, respectively. The excess heparinase I was inactivated once the increase reached the endpoint by heating the solution at 100 °C for 10 min. The solution was filtered, precipitated with alcohol, and lyophilized to obtain NALHP and NAULHP. The HP derivative NAI45 was prepared through the enzymolysis of HP using the heparinase I, allowing the increase in the absorbance to reach 45, followed by periodate sodium oxidation and sodium borohydride reduction. The deanticoagulated enoxaparin (NAEno) was prepared from commercial enoxaparin (Changshan Biochemical Pharmaceutical Co., Ltd., Hebei, China) using the same preparation process used for the oxidation of HP to NAHP.

### Molecular Weight Determination of Deanticoagulated HPs

The molecular weight of each HP derivatives was determined using gel penetration chromatography (GPC) as described previously. Briefly, an HPLC system (Waters, USA) equipped with a binary pump module, an autosampler, and a refractive index detector was used. The mobile phase consisted of 0.2 M sodium sulfate solution at pH 5.0. HP derivatives, the HP sodium molecular weight calibrant (USP, USA), and low-molecular-weight HP for molecular weight calibration (NIBSC, USA) were dissolved in the mobile phase at a concentration of 20 mg/ml. The samples were subsequently injected onto a 300 × 7.8-mm TSK Gel G2000SW_XL_ column (Tosoh, Japan) in a volume of 25 μl. The column was eluted at a flow rate of 0.5 ml/min and a temperature of 35 °C. The data were collected and analyzed in Breeze 2 (Waters, USA) software. The molecular weight distributions of the HP derivatives were calculated based on the standard curve generated from the calibrant with a general third-order calibration method.

### Fine separation of NALHP

Fine separation of the HP derivatives was performed using a HiPrep 16/60 Sephacryl S-100 HR column (GE Healthcare, USA) with a ÄKTA prime purification system (GE Healthcare, USA). The mobile phase consisted of 0.2 M sodium chloride solution at pH 5.0. The HP derivatives NALHP was dissolved in the mobile phase at a concentration of 120 mg/ml. The sample was subsequently injected onto the column using a 5 ml injection loop. The column was eluted at a flow rate of 0.5 ml/min at room temperature. Fractions were collected every 10 min from 70 min to 180 min after sample injection and were designated S1 to S11 according to the order of retention time. Afterward, the samples were lyophilized and reconstituted in water at 30% of the original volume to enrich polysaccharides, and alcohol precipitation and another lyophilization step were then performed to obtain the separated components.

### Measurement of residual anti-factor Xa activity

Residual anticoagulant activity in heparin, NAHP, NALHP and S6 was evaluated using an antithrombin III-dependent chromogenic anti-factor Xa assay, adapted from the European Pharmacopoeia method for low-molecular-weight heparins.

Samples and the low-molecular-weight heparin biological reference standard were diluted in Tris-HCl buffer (pH 7.4) to generate four concentrations within the linear range of the dose–response relationship. Adjacent concentrations were prepared at a dilution ratio of approximately 1:0.6– 1:0.7, and each concentration was tested in duplicate.

Sample or reference-standard solution was incubated with antithrombin III, followed by the addition of factor Xa. Residual factor Xa activity was determined using the chromogenic substrate S-2765. After incubation at 37°C, the reaction was terminated with 30% acetic acid, and absorbance was measured at 405 nm. Anti-Xa activity was calculated relative to the reference standard using a 4×4 parallel-line bioassay model and expressed as international units per milligram of sample (IU/mg).

Anti-Xa activity was used to determine whether the periodate oxidation–borohydride reduction and subsequent enzymatic depolymerization and fractionation steps effectively reduced the anticoagulant activity of NAHP, NALHP and S6.

### Mouse model of DSS-induced colitis

Male C57BL/6J mice (18–22 g, 6–8 weeks) were purchased from Vital River Laboratories (Beijing, China). The mice were housed under specific pathogen-free conditions in the laboratory animal facility of Tsinghua University. The animal protocols 17-XXH1 and 20-XXH1 were carried out with the approval of the Institutional Animal Care and Use Committee (IACUC) of Tsinghua University. The laboratory animal facility was accredited by AAALAC (Association for Assessment and Accreditation of Laboratory Animal Care) International.

Evaluation of the anti-UC efficacy of the HP derivatives was performed using a 7-day mouse model of acute colitis. All mice were pre-fed for one week prior to any subsequent experiment. The mice were given 3% DSS (dextran sulfate sodium; colitis grade, MP Biomedicals, USA) solution instead of water for 7 consecutive days. The HP derivatives and 5-aminosalicylic acid (5-ASA) were dissolved in saline at a concentration of 3 mg/ml. Starting 3 days post-DSS induction, the mice were orally administered the drug solution daily at a dosage of 30 mg/kg/d. The mice were euthanized by CO_2_ inhalation after DSS induction for 7 days. The serum was separated from the blood of the mice obtained by cardiac puncture, whereas the spleens and the colons were excised and measured. The colon tissue was cut in the longitudinal direction, washed with ice-cold PBS, rolled with the Swiss-rolling technique, fixed, and embedded in paraffin for further investigations.

Multiomics analyses were performed in a 14-day mouse colitis model to evaluate the systemic changes in mice with colitis caused by drugs. The mice were allowed to drink 3% DSS solution for 3 days, after which they received standard drinking water and drug treatment for 4 days. Afterward, the mice were allowed to drink 2% DSS solution for another 3 days, after which they received standard drinking water and drug treatment for another 4 days. Fresh feces were collected prior to euthanasia by CO_2_ inhalation. After the mice were sacrificed, the colons were removed. The colonic contents were collected while the colonic epithelial tissue was scraped and collected in 1.5 ml centrifuge tubes containing TRIzol reagent (Invitrogen, USA).

### ELISA

The concentrations of interleukin-6 (IL-6) and tumor necrosis factor-alpha (TNF-α) in serum and cell culture medium were measured using an IL-6 Mouse ELISA Kit (Invitrogen, USA) and a TNF-α Mouse ELISA Kit (Invitrogen, USA), respectively, according to the manufacturer’s instructions.

### Biolayer interferometry

The molecular interactions between DSS and the HP derivatives were evaluated using an Octet Red96 instrument (FortéBIO, USA). DSS and the HP derivatives were dissolved in HBS-EP buffer containing 10 mM HEPES, 150 mM NaCl, 3 mM EDTA, and 0.005% Tween 20 at pH 7.4. The streptavidin sensors were loaded with DSS washed in HEB-EP buffer and exposed to the HP derivatives.

### RNA isolation and qPCR

Total RNA was extracted using TRIzol reagent (Invitrogen) and reverse transcribed to cDNA using a FastKing RT Kit (TIANGEN, China). All qPCRs were run in a CFX96 Touch Real-Time PCR Detection System (Bio-Rad, USA) using Talent qPCR PreMix (TIANGEN, China) with the specific primers listed in the supplementary materials.

### FTIR spectroscopy

The infrared absorption spectra of the HP derivatives were generated using a Nicolet 6700 FTIR spectrometer (Thermo Electron Corporation, Shanghai, China) with the potassium bromide tablet method. The spectral data were collected in a 4000–400 cm^-1^ range for all the samples.

### NMR analysis

For 1D (^1^H, ^13^C) and 2D-NMR analyses, HP derivatives (50–60 mg) were dissolved in 0.6 mL of deuterium oxide and lyophilized repeatedly to remove exchangeable protons. The thoroughly dried samples were then dissolved in 0.5 mL of deuterium oxide with 0.002% 3-(trimethylsilyl) propionate sodium salt (TSP) and transferred into 25 × 0.5-cm NMR tubes (Wilmad Lab Glass, USA). 1D (^1^H, ^13^C) and 2D-NMR experiments were performed at 298 K on a Bruker Advance 600 MHz spectrometer (Germany) equipped with a 5 mm helium-cooled cryoprobe using the following parameters: Proton—number of scans, 16; number of dummy scans, 2; relaxation delay, 1.0 s; spectral width, 20 ppm; and transmitter offset, 6.0 ppm. One-dimensional ^13^C spectra were obtained with proton decoupling during acquisition at 298 K over 5000 scans. All the spectra were Fourier transformed, phased and baseline corrected. The chemical shift values were calibrated with TSP at 0.00 ppm. HSQC spectra were recorded with carbon decoupling during acquisition with 256 increments of 16 scans. The polarization transfer delay was set for a ^1^J_C-H_ coupling value of 145 Hz.

### Microbiome analysis

#### Sample preparation and DNA extraction

Fresh fecal samples from UC mice were collected in 1.5-ml centrifuge tubes and stored immediately at –80 °C until further analysis. Microbial genomic DNA samples were extracted using a DNeasy PowerSoil Kit (QIAGEN, Germany) following the manufacturer’s instructions. The DNA concentrations in the samples were subsequently quantified using a NanoDrop ND-1000 spectrophotometer (Thermo Fisher Scientific, USA), and the samples were then qualified by agarose gel electrophoresis.

#### 16S rRNA amplicon sequencing

The V3–V4 region of the bacterial 16S rRNA gene was amplified by PCR using the forward primer 338F (5’-ACTCCTACGGGAGGCAGCA-3’) and the reverse primer 806R (5’-CGGACTACHVGGGTWTCTAAT-3’). Sample-specific 7-bp barcodes were added to the primers for further multiplex sequencing. PCR amplicons were purified using VAHTS^TM^ DNA Clean Beads (Vazyme, China) and quantified using a PicoGreen dsDNA Assay Kit (Invitrogen, USA) prior to manipulation to achieve equal amounts of DNA. Afterward, paired-end 2 × 250 bp sequencing was performed on the MiSeq platform (Illumina, USA) with a NovaSeq 6000 SP Reagent Kit.

#### Microbiome data analysis

The raw sequence data were processed with QIIME2 2019.4 (Bolyen *et al*. 2018) with slight modifications based on the official tutorials (https://docs.qiime2.org/2019.4/tutorials/). In brief, the raw data were demultiplexed with the demux plugin, and primers were removed with the cut adapt plugin. Afterward, the data were subjected to quality filtering, denoising, merging, and chimera removal with the DADA2 plugin. A count table without rarefaction was used for further analysis. The following analysis was carried out using an in-house-developed R script in RStudio software (v.2021.09.1, R version 4.1.2). Alpha diversity analysis and principal coordinate analysis (PCoA) were performed using the vegan (v.2.5-7) package. Enterotyping was performed with the Dirichlet multinomial mixture model with the DirichletMultinomial (v.1.36.0) package. Data visualization was performed using the ggplot2 (v.3.3.5) and ggalluvial (v.0.12.3) packages. In addition, LEFSe analysis was performed on the Genescloud platform (https://www.genescloud.cn/home).

### Metabolome analysis

#### Sample preparation and extraction

Samples of the colonic contents were thawed on ice. Aqueous homogenates were obtained by mixing 100 mg of each sample with 300 μl of ice-cold water and steel balls for 5 min prior to incubation on ice for 10 min. Then, 500 μl of methanol was added, and the homogenates were mixed by vortexing for 5 min prior to incubation on ice for another 10 min. The homogenates were centrifuged (12,000 rpm, 4 °C, 10 min), and 600 μl of the supernatant was transferred to 1.5-ml centrifuge tubes for further concentration. The concentrated dry products were dissolved in 100 μl of 5% methanol–water, vortexed, and centrifuged (12,000 rpm, 4 °C, 10 min). Finally, the supernatants were collected for LC‒MS/MS analysis.

#### LC‒MS analysis

A combination of positive ion mode MS and negative ion mode MS was applied to identify the metabolites in the colonic contents. Data were acquired with an LC‒MS system comprising a 1290 Infinity LC System (Agilent, USA) coupled to a 6545 LC/Q-TOF mass spectrometer (Agilent, USA). The method details are summarized as follows. The supernatants were injected onto a 100 × 2.1-mm ACQUITY UPLC HSS T3 C18 column (Waters, USA) in a volume of 2 μl. The column was eluted at a flow rate of 400 μl/min and a temperature of 40 °C. Mobile phase A (water with 0.1% formic acid) and mobile phase B (acetonitrile with 0.1% formic acid) were used with the following gradient program: 95:5 V/V A:B at 0 min, 10:90 V/V at 11.0 min, 10:90 V/V at 12.0 min, 95:5 V/V at 12.1 min, 95:5 V/V at 14.0 min. In positive ion mode, MS analysis was carried out using the following settings: ion spray voltage, 250; gas flow, 8; fragmentor, 135; gas temperature, 325 °C; sheath temperature, 325 °C; sheath flow, 11; and nebulizer, 40. In negative ion mode, MS analysis was carried out using the following settings: ion spray voltage, 1,500; gas flow, 8; fragmentor, 135; gas temperature, 325 °C; sheath temperature, 325 °C; sheath flow, 11; nebulizer, 40.

#### Metabolome data analysis

The raw data obtained by LC‒MS analysis were first converted into mzML format with ProteoWizard software. Peak extraction, alignment, and retention time correction were carried out with the XCMS program, while the peak area was corrected using the “SVR” method. Peaks with a deletion rate of > 50% were filtered out in each group of samples. Next, metabolic identification information was obtained by searching the laboratory’s in-house-built database and integrating the public database and the metDNA tool. Finally, statistical analysis was carried out using an in-house-developed R script in RStudio software (v.2021.09.1, R version 4.1.2). Metabolites at the MS2 confidence level were selected for the following analysis. Principal component analysis (PCA) was performed and the results were visualized using the vegan (v.2.5–7) and ggplot2 (v.3.3.5) packages, respectively. Orthogonal partial least squares discriminant analysis (OPLS-DA) was carried out using the ropls (v.1.26.0) package, and metabolites with a VIP > 1, p < 0.05, and |logFC| > 1 were identified as differentially abundant metabolites. The abundances of the differentially abundant metabolites in each group were visualized using the ComplexHeatmap (v.2.10.0) package. Spearman correlation analysis was performed on the the metabolome and microbiome data, and the results were visualized using the WGCNA (v.1.70-3) and pheatmap (v.1.0.12) packages, respectively. The Mantel test was performed between the metabolome and microbiome data using the ggcor (v.0.9.4.3) package.

### Transcriptome analysis

#### RNA-seq library construction and sequencing

Total RNA from the colonic epithelial tissue of UC mice was extracted using TRIzol reagent (Invitrogen, USA). The RNA-seq libraries were constructed using a TruSeq RNA Sample Prep Kit (Illumina, USA) according to the manufacturer’s instructions. The polyA+ RNA was enriched, fragmented, and reverse transcribed to obtain cDNA, and end repair, PCR amplification, purification, and quantification were then performed. The libraries were subsequently sequenced on the Illumina NovaSeq 6000 platform.

#### RNA-seq data analysis

The raw data were quality filtered using fastx_toolkit (v.0.0.14), mapped to the mouse genome GRCm39 using Bowtie2 (v.2.4.1), assembled with cufflinks (v.2.2.1), and quantified with RSEM (v.1.1.3). The transcripts per million (TPM) values were used in the following analysis. Low-abundance data were filtered based on the mean TPM values. The differentially expressed genes were defined as those whose p value was <0.05 with the DESeq2 (v.1.34.0) package, and GO and KEGG enrichment analyses were then performed with the clusterProfiler (v.4.2.1) package. The results were visualized using the ggplot2 (v.3.3.5) and ComplexHeatmap (v.2.10.0) packages.

### Cell culture

All the cells were maintained at 37 °C in 5% CO_2_ in a Heracell 150i incubator (Thermo Fisher Scientific, USA). Human NCM460 cells (ATCC) were cultured in RPMI 1640 medium (Gibco, USA) supplemented with 10% fetal bovine serum (Gemini, USA). Mouse RAW264.7 cells (ATCC) were cultured in DMEM (Gibco, USA) supplemented with 10% fetal bovine serum.

To assess the anti-inflammatory effects of the HP derivatives, RAW264.7 cells were seeded in 96-well plates, stimulated with 100 ng/ml lipopolysaccharide (Sigma‒Aldrich, USA) for 15 min, and treated with the HP derivatives at a concentration of 1 mg/ml for 24 hr. The supernatants were collected and subjected to ELISA.

### UC organoid culture

Human UC patient-derived colonoids were cultured as previously described ^35,36^ with slight modifications. Colon tissue samples were obtained from biopsies taken from male and female UC patients aged 20 to 80 years (inclusive); in all patients, UC had been clearly diagnosed for more than 3 months through clinical manifestations and endoscopic examination. All samples were graded as mild to moderate active ulcerative colitis (Mayo score: 3–10) and were collected at Tsinghua Changgung Hospital in Beijing, washed in ice-cold PBS, incubated with 10 mM DTT for 15 min, and disassociated in PBS containing 20 mM EDTA for 60 min at 4 °C. The crypts were isolated by vortexing, collected by centrifugation, embedded in growth factor-reduced Matrigel (Corning), and cultured in complete medium consisting of advanced DMEM/F12 (Thermo Fisher Scientific, USA) supplemented with 1% penicillin and streptomycin (Thermo Fisher Scientific, USA), 10 mM HEPES (Thermo Fisher Scientific, USA), 2 mM GlutaMAX (Thermo Fisher Scientific, USA), 1 × B-27 supplement (Thermo Fisher Scientific, USA), 1 mM N-acetylcysteine (Sigma‒Aldrich, USA), 100 ng/ml recombinant mouse EGF (Thermo Fisher Scientific, USA), 100 ng/ml recombinant human Noggin (PeproTech, USA), 1 μg/ml recombinant human R-spondin-1 (PeproTech, USA), 500 nM A83-01 (Tocris, England), 10 μM SB202190 (Sigma‒Aldrich, USA) and 10% Afamin/WNT3A conditioned medium (JSR Life Sciences, Japan). To prevent anoikis, 10 mM Y-27632 (STEMCELL Technologies, Canada) was added to the complete medium for the first day of culture. The colonoids were passaged every 6 to 7 days by mechanical disassociation, and the medium was changed every day to ensure their rapid growth.

Human samples were collected under protocols approved by the Ethics Committee of Beijing Tsinghua Changgung Hospital (Approval No. 22111–4-02). Written informed consent was obtained from all the participants.

### Western blotting and nuclear/cytoplasmic fractionation

Organoids were collected after the indicated treatments, washed with ice-cold PBS and lysed for total-protein extraction or nuclear/cytoplasmic fractionation. Nuclear and cytoplasmic proteins were isolated using a Nuclear and Cytoplasmic Protein Extraction Kit (Beyotime, China). Protein concentrations were determined by BCA assay. Equal amounts of protein were separated by SDS-PAGE and transferred to PVDF membranes. Membranes were blocked with 5% bovine serum albumin in TBST for 1 hour at room temperature and incubated overnight at 4 C with primary antibodies against MUC2, CTNNB1, TCF1/7, HES1, GAPDH and LAMINB. After washing, membranes were incubated with HRP-conjugated secondary antibodies for 1 hour at room temperature. Signals were detected using Omni-ECL Femto Light Chemiluminescence Kit (Epizyme Biotech, China) and imaged using a ChemiDoc system (Bio-Rad, USA). Band intensities were quantified using ImageJ; cytoplasmic proteins were normalized to GAPDH and nuclear proteins to LAMINB.

### EdU labeling

EdU labeling of proliferating cells in the colonic epithelia of UC mice was performed on 3-μm sections of paraffin-embedded tissues with a Click-iT EdU Imaging Kit (Invitrogen, USA) according to the manufacturer’s protocol.

### Immunohistochemical and immunofluorescence staining

Paraffin-embedded sections (3 μm thick) were deparaffinized, washed with PBS containing 0.01% Tween 20, and subjected to antigen retrieval by immersion in Tris-EDTA buffer (pH 9.0) at 100 °C for 20 min.

For immunohistochemical staining, the samples were blocked using peroxidase-blocking solution (Dako, Denmark) and incubated with specific primary antibodies at 4 °C overnight. The fluorescence signals were detected using a REAL EnVision Detection System (Dako, Denmark) and observed using Axioscan. Z1 Slide Scanner (Zeiss, Germany).

For immunofluorescence staining, the samples were blocked with 1% BSA/PBS for 1 hr, incubated with specific primary antibodies at 4 °C overnight, and then incubated with secondary antibodies; nuclei were then stained, and the samples were mounted with ProLong Gold Antifade Mountant (Thermo Fisher Scientific, USA) and observed using inverted confocal microscopy (Nikon, Japan).

### Single-cell RNA sequencing ^33^

#### Preparation of single-cell suspensions from UC organoids

The UC organoids grown on filter paper were placed on an ice-cold plate and allowed to cool for 2–3 minutes. The sample wells containing the tissue spheroids to be analyzed were confirmed, the culture medium in the wells was discarded, and the solid Matrigel^TM^ was left. One milliliter of precooled (4 °C) advanced DMEM/F12 culture medium supplemented with 1% HEPES, 1% P/S and 1% GlutaMAX was added to each well, the Matrigel^TM^ surrounding the UC organoids was gently disrupted by pipetting to break it down and dissolve it, and the organoids were collected in 15 mL centrifuge tubes. This procedure was repeated 3 times until all the tissue spheroids were disrupted by pipetting and collected. The mixture was centrifuged at 300 × g and 4 °C for 5 minutes, the supernatant was removed, 3 mL of precooled (4 °C) advanced DMEM/F12 culture medium was added to resuspend the pellet, and the mixture was centrifuged again at 200 × g and 4 °C for 5 minutes to remove the supernatant to obtain the colon tissue spheroids. Two milliliters of 0.25% trypsin was added to the suspension, which was subsequently incubated at 37 °C in a water bath for 5 minutes, after which an equal volume of RPMI-1640 medium containing 10% fetal bovine serum (FBS) was added to terminate the enzymatic digestion. A 1 mL pipet was used to pipet the suspension ten times. The suspension was centrifuged at 300 × g and 4 °C for 5 minutes, and the supernatant was removed. Then, 2 mL of 0.0005% Y-27632/TrypLE was added to generate a suspension, which was incubated at 37 °C in a water bath for 15 minutes and pipetted ten times at 5-minute intervals. After the incubation was complete, an equal volume of RPMI-1640 medium containing 10% FBS was added to terminate the reaction. The suspension was centrifuged at 400 × g and 4 °C for 5 minutes, and the supernatant was removed. Then, 50–200 μL of RPMI-1640 medium containing 10% FBS was added to generate a suspension so that the cell concentration was within the range of 700–1200 cells/μL. This suspension was disrupted by vigorous pipetting ten to twenty times to obtain single cells from the UC organoids.

#### Single-cell RNA sequencing and library construction for colonic organoids

The concentration and viability of the cells in the single-cell suspensions of colonic organoids were measured with a cell counter. Samples with a cell viability greater than 80%, an agglomeration rate of less than 5% and a lack of obvious cell debris or impurities were considered suitable for library construction. Using microfluidic technology, gel beads and individual cells with specific cell barcodes and reverse transcription information were coated with oil droplets. The cells were solubilized and released mRNA in this system, and the mRNA was captured by the gel beads and reverse transcribed to generate the full-length cDNA sequence. The oil droplets were then disrupted and purified, the amount of cDNA was increased by PCR amplification, and the sequencing primers were ligated. The cDNA was cleaved into fragments of approximately 200∼300 bp by a chemical method, end repair was carried out, P7 adapters were connected, sample barcodes were introduced, and the sequence fragments were screened. The library was sequenced on the Illumina NovaSeq platform with a PE150 sequencing strategy.

The unique molecular identifiers (UMIs) in the off-machine data were preprocessed using CellRanger software, an official tool of 10x Genomics. Loupe Browser 6 software, an official tool of 10x Genomics, was used to delete the multicell data, low-quality data, and apoptotic cell data. The corresponding data barcode criteria were a UMI count of greater than 60,000, a feature count of less than 210, and a percentage of mitochondrial UMIs of greater than 25%. Afterward, the uniform manifold approximation and projection (UMAP) method was used for dimensionality reduction. After dimensionality reduction, the CellRanger off-machine data were imported into R Studio, and Seurat was used for dimensionality reduction classification.

## Data availability

All the data supporting the results of this study are available within the paper and its Supplementary Materials. Additional processed data are available from the corresponding authors upon request.

## Acknowledgements

We thank the Biopharmaceutical and Health Engineering Research Core Facility, Tsinghua Shenzhen International Graduate School, Tsinghua University, for technical assistance; the National Center for Electron Microscopy in Beijing, School of Materials Science, Tsinghua University, for assistance with electron microscopy imaging; the Laboratory Animal Resources Center, Tsinghua University, for technical support.

## Contributors

Conceptualization: XH. X., Y. W. and X. J.

Methodology: W. H., L. H., Y. W., W. Z., Y. J., ZY. S., HH. C. and XH. X.

Visualization: W. H., L. H. and W. Z.

Investigation: W. H., L. H., ZK. L., QX. Z., W. Z. and XY. R.

Validation: W. H., Y. W. and ZQ. Z.

Funding acquisition: XH. X., Y. W. and X. J.

Supervision: XH. X., Y. W. and X. J.

Writing – original draft: W. H., Y. W. and XH. X.

Writing – review and editing: W. H., Y. W., XH. X., YS. Y., HC. Z., CY. Z., Y. C., PE. L., B. X. and C. Z.

All authors reviewed and approved the final version of the manuscript. XH. X. is the guarantor of the study and accepts full responsibility for the conduct of the study, had access to the data, and controlled the decision to publish.

## Ethics approval

Human colon tissue samples were collected under protocols approved by the Ethics Committee of Beijing Tsinghua Changgung Hospital (approval No. 22111-4-02). Written informed consent was obtained from all participants before sample collection. The study was conducted in accordance with the Declaration of Helsinki.

All animal procedures were approved by the Institutional Animal Care and Use Committee of Tsinghua University under protocols 17-XXH1 and 20-XXH1. The animal facility is accredited by AAALAC International, and the animal experiments were reported in accordance with the ARRIVE guidelines where applicable.

## Data availability statement

RNA-sequencing, single-cell RNA-sequencing, metabolomics and 16S rRNA sequencing data generated in this study have been deposited. Source data underlying the quantitative graphs and uncropped immunoblots are provided as source data files. Additional processed data supporting the findings of this study are available from the corresponding author upon reasonable request.

Custom R scripts used for the transcriptomic, microbiome, metabolomic and single-cell analyses are available through GitHub or from the corresponding author upon reasonable request. NALHP, S6 and related deanticoagulated HPs generated in this study are available from the corresponding author upon reasonable request and subject to institutional material transfer agreements.

## Supplementary Materials

**Supplementary Fig. 1.**
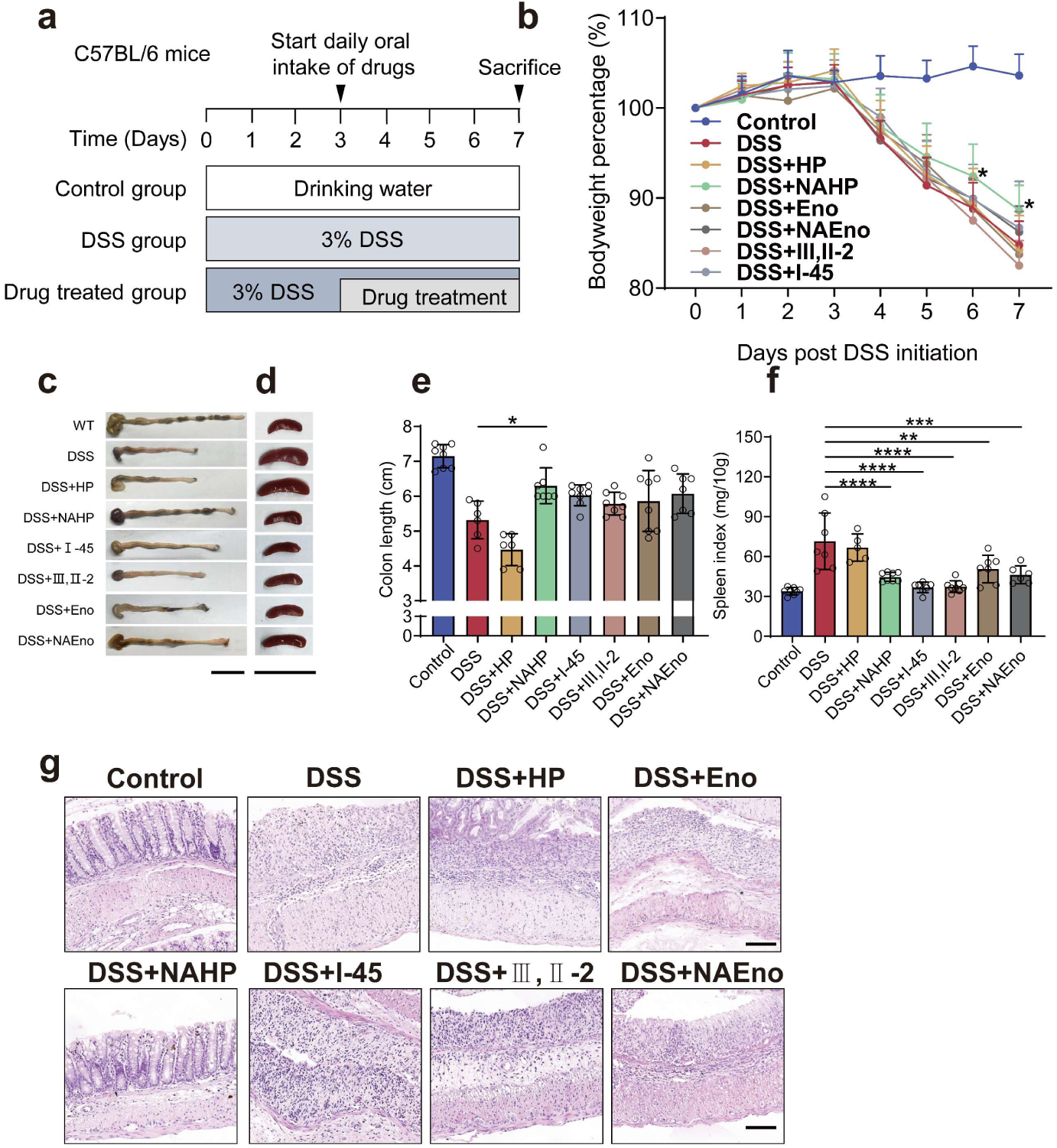
Activity-guided screening identifies anti-colitis activity among deanticoagulated heparin derivatives. **a**, Experimental design for screening the deanticoagulated heparin library in DSS-induced acute colitis. Mice received 3% DSS for 7 days, and oral treatment with native heparin (HP), enoxaparin (Eno), non-anticoagulant heparin (NAHP), non-anticoagulant enoxaparin (NAEno), or enzymatically depolymerized heparin derivatives was initiated on day 3. **b**, Changes in body weight during DSS exposure and treatment. **c-d**, Representative gross images of colons (**c**) and spleens (**d**) collected at the experimental endpoint. **e-f**, Quantification of colon length (**e**) and spleen index (**f**). **g**, Representative H&E-stained colon sections showing differential recovery of mucosal architecture among heparin derivatives. Scale bar, 100 μm. The data are presented as the mean ± SD. * *p* < 0.05, ** *p* < 0.01, *** *p* < 0.001, **** *p* < 0.0001; ns (not significant) *p* > 0.05; analyzed by ordinary one-way ANOVA multiple comparisons test in (**b**).

**Supplementary Fig. 2.**
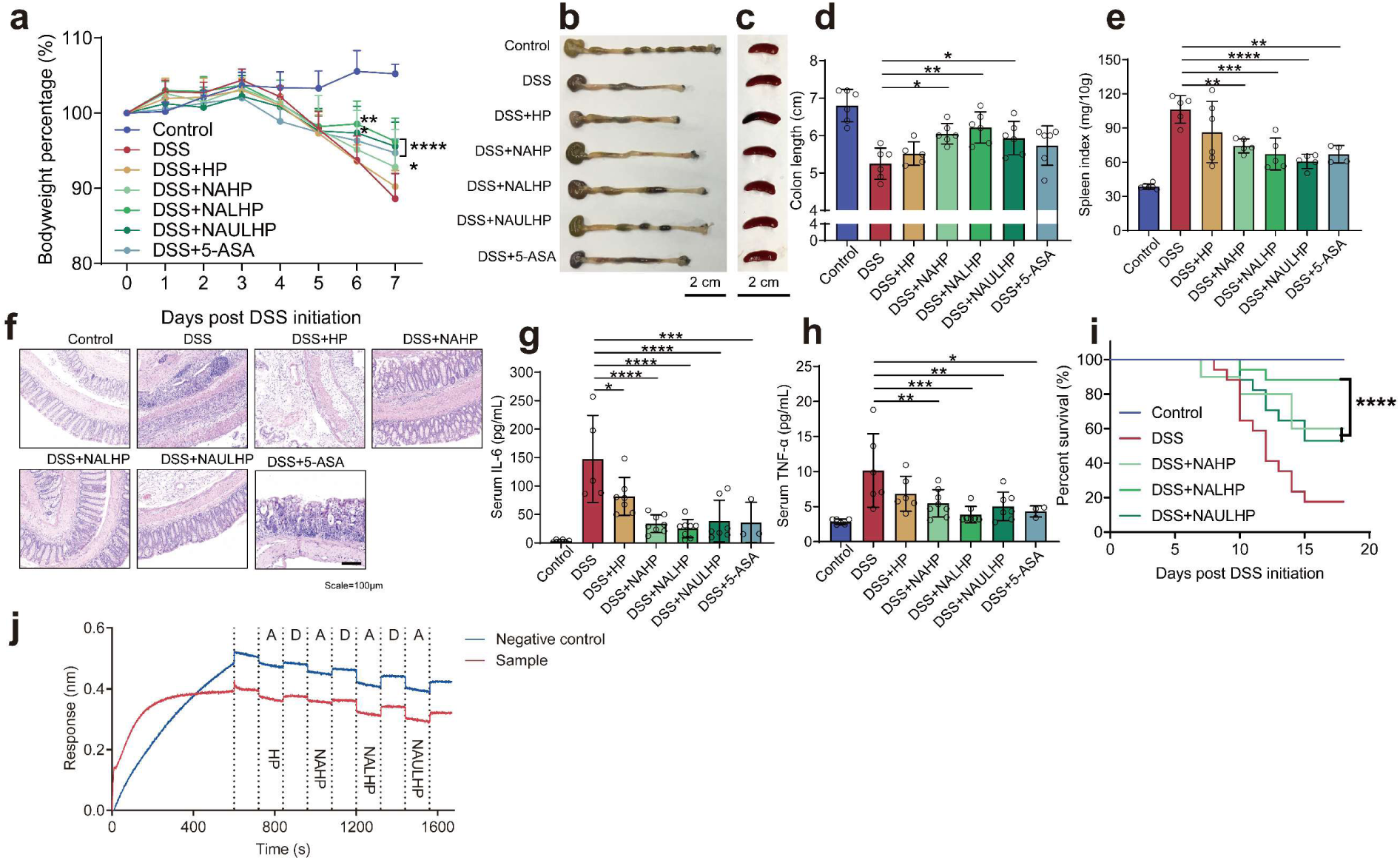
Molecular-weight reduction enriches the therapeutic activity of non-anticoagulant heparin. **a**, Changes in body weight in DSS-treated mice receiving HP, NAHP, NALHP, NAULHP or 5-ASA. **b-c**, Representative gross images of colons (**b**) and spleens (**c**). **d-e**, Colon length (**d**) and spleen index (**e**). **f**, Representative H&E-stained colon sections. Scale bar, 100 μm. **g-h**, Serum IL-6 (**g**) and TNF-α (**h**) concentrations. **i**, Survival curves of DSS-treated mice receiving NAHP derivatives with different molecular-weight distributions. **j**, Representative BLI response curves comparing interactions associated with HP, NAHP, NALHP and NAULHP preparations. The blue line represents the blank control, and the orange line represents the sample. (**a, d, e, g-i**) Representative images or quantitative analysis from at least n=5 biologically independent samples are shown. The data are presented as the mean ±SD. * *p* < 0.05, ** *p* < 0.01, *** *p* < 0.001, **** *p* < 0.0001, and ns (not significant) *p* > 0.05, analyzed by ordinary one-way ANOVA multiple comparisons test in (**A**).

**Supplementary Fig. 3.**
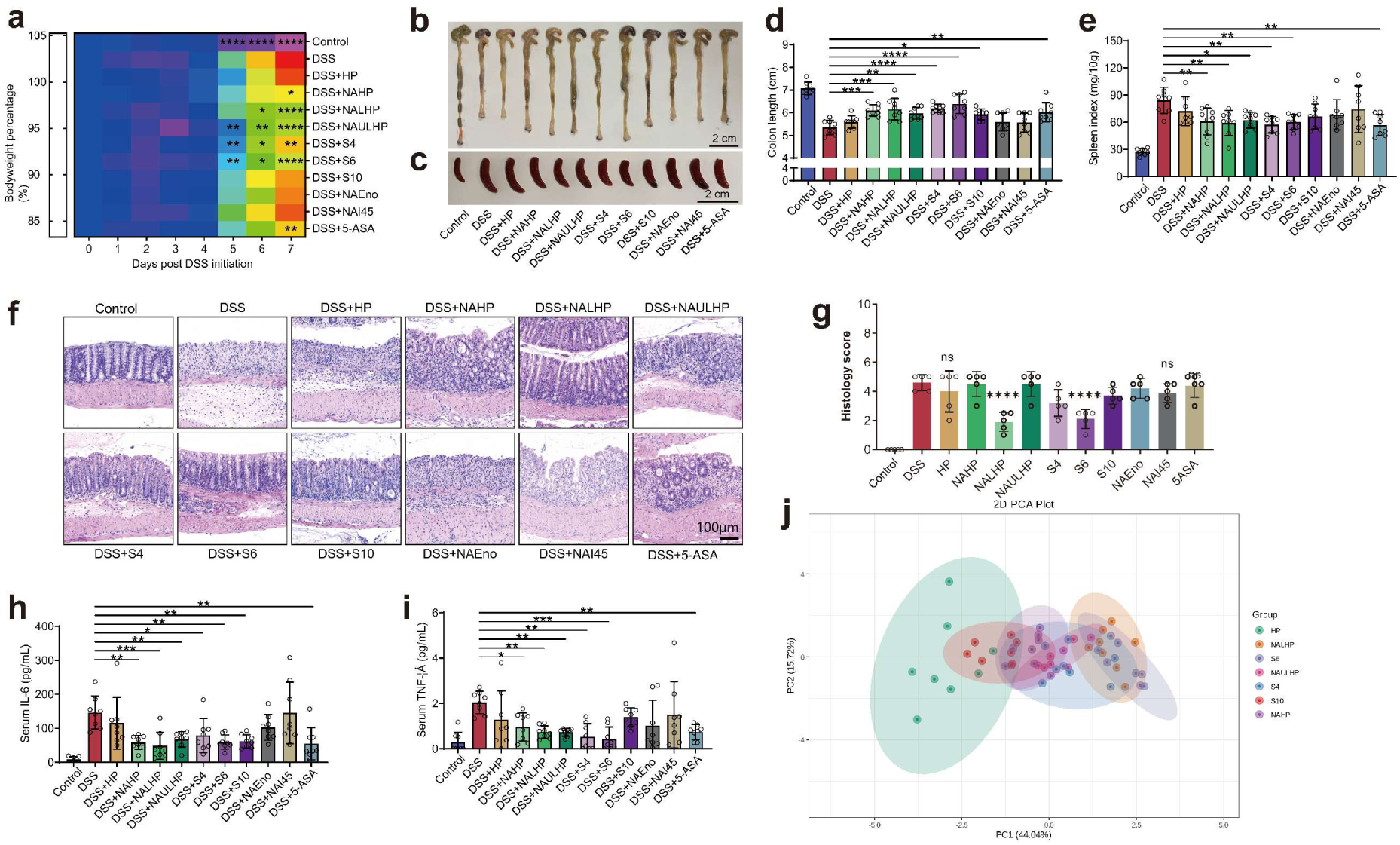
Therapeutic efficacy is enriched in defined non-anticoagulant low-molecular-weight heparin fractions. **a**, Heatmap of body-weight responses in DSS-treated mice receiving the indicated native, deanticoagulated or fractionated heparin preparations. **b-c**, Representative gross images of colons (**b**) and spleens (**c**). **d-e**, Colon length (**d**) and spleen index (**e**). **f-g**, Representative H&E staining (**f**) and histological injury scores (**g**). Scale bar, 100 μm. **h-i**, Serum IL-6 (**h**) and TNF-α (**i**) concentrations. **j**, Principal-component analysis integrating body weight, colon length, spleen index, inflammatory cytokines and histological scores across treatments. (**d, e, g-i**) Representative images or quantitative analysis from n=6 biologically independent samples are shown. The data are presented as the mean ±SD. * *p* < 0.05, ** *p* < 0.01, *** *p* < 0.001, **** *p* < 0.0001, and ns (not significant) *p* > 0.05, analyzed by ordinary one-way ANOVA multiple comparisons test in (**a**). The PCA data represent the improvement in the characteristics of colitis mice (including body weight, colon length, spleen index, serum IL-6 and TNF-α levels, and histological scores) treated with drugs of different molecular weights (HP, NAHP, NALHP, NAULHP, S4, S6 and S10).

**Supplementary Fig. 4.**
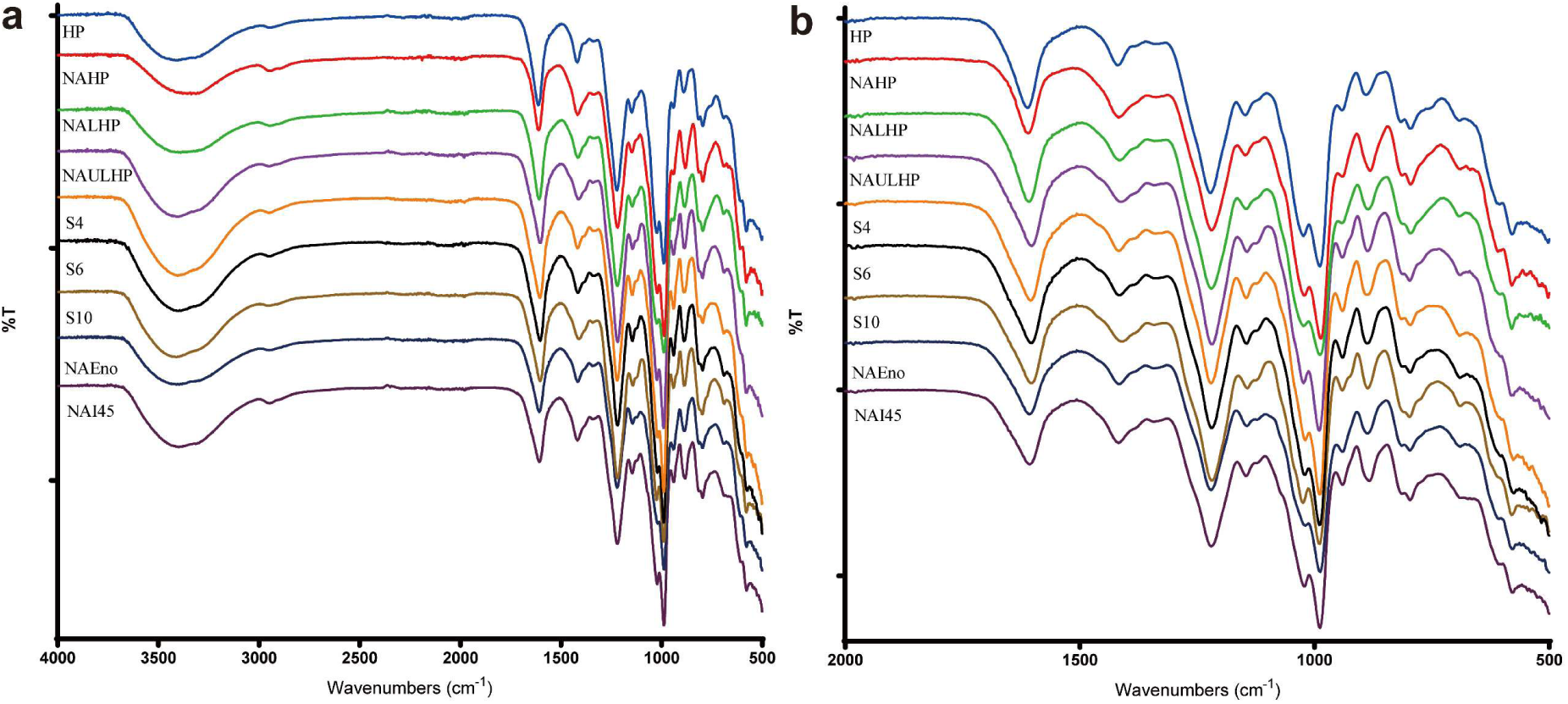
Infrared spectroscopy confirms conserved heparin-like structural features after deanticoagulation and fractionation. **a**, Full FTIR spectra of native and modified heparin preparations. **b**, Enlarged spectra between 2000 and 500 cm⁻¹ highlighting characteristic glycosidic, carboxylate and sulfate-associated absorption regions.

**Supplementary Fig. 5.**
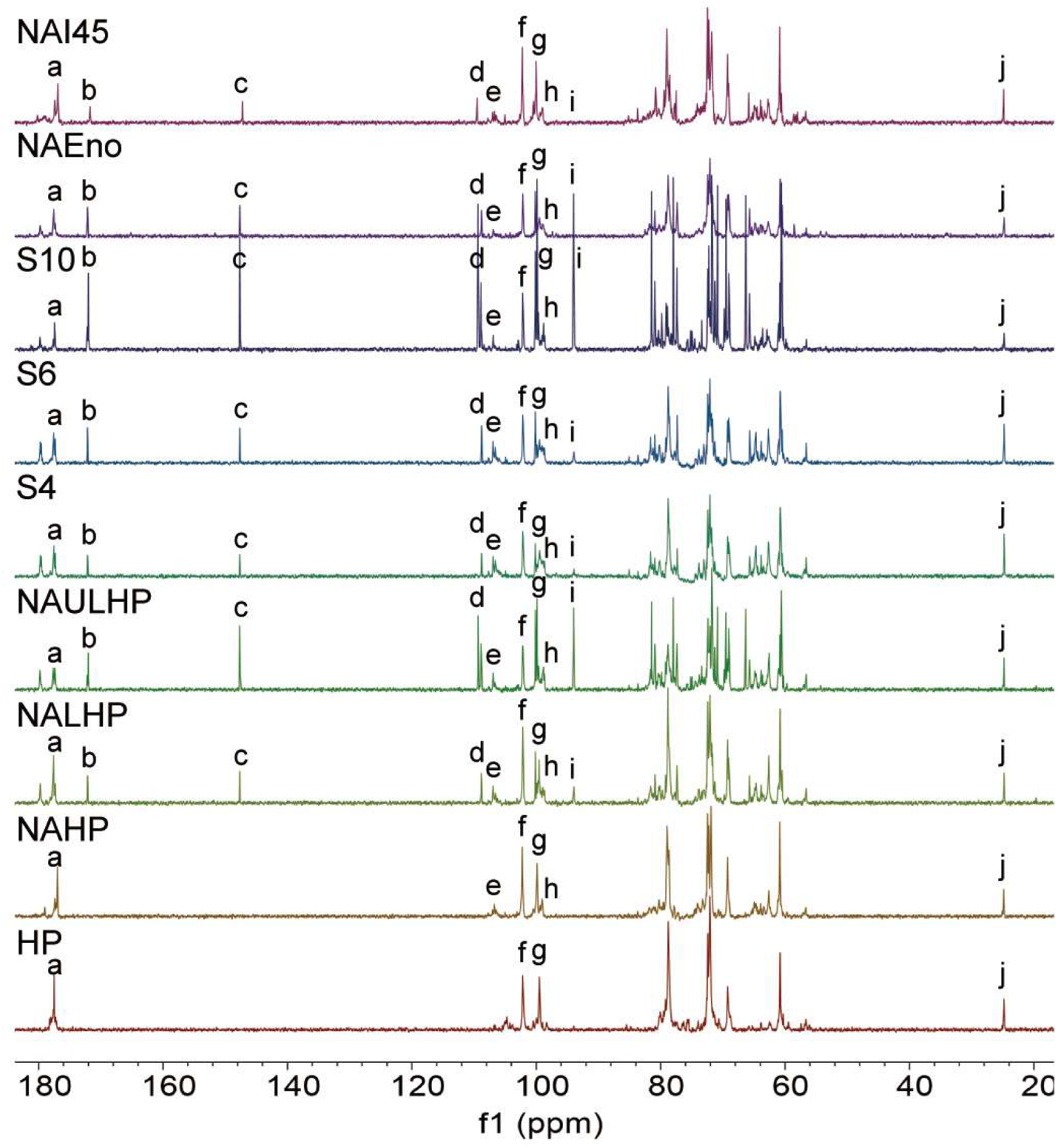
^13^C NMR spectra characterize structural features of native and modified heparin preparations.

**Supplementary Fig. 6.**
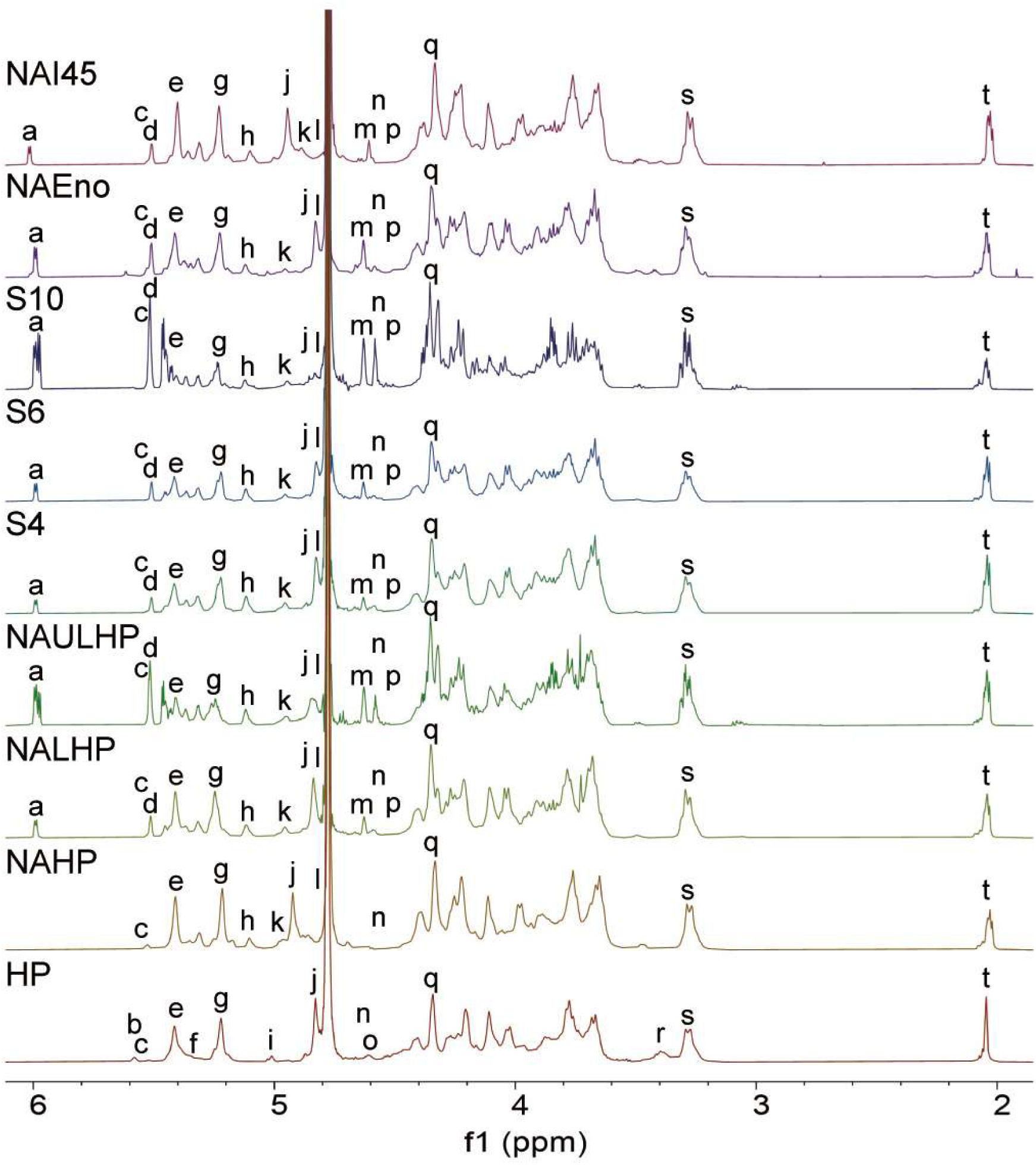
^1^H NMR spectra reveal conserved and differential structural features among heparin derivatives.

**Supplementary Fig. 7.**
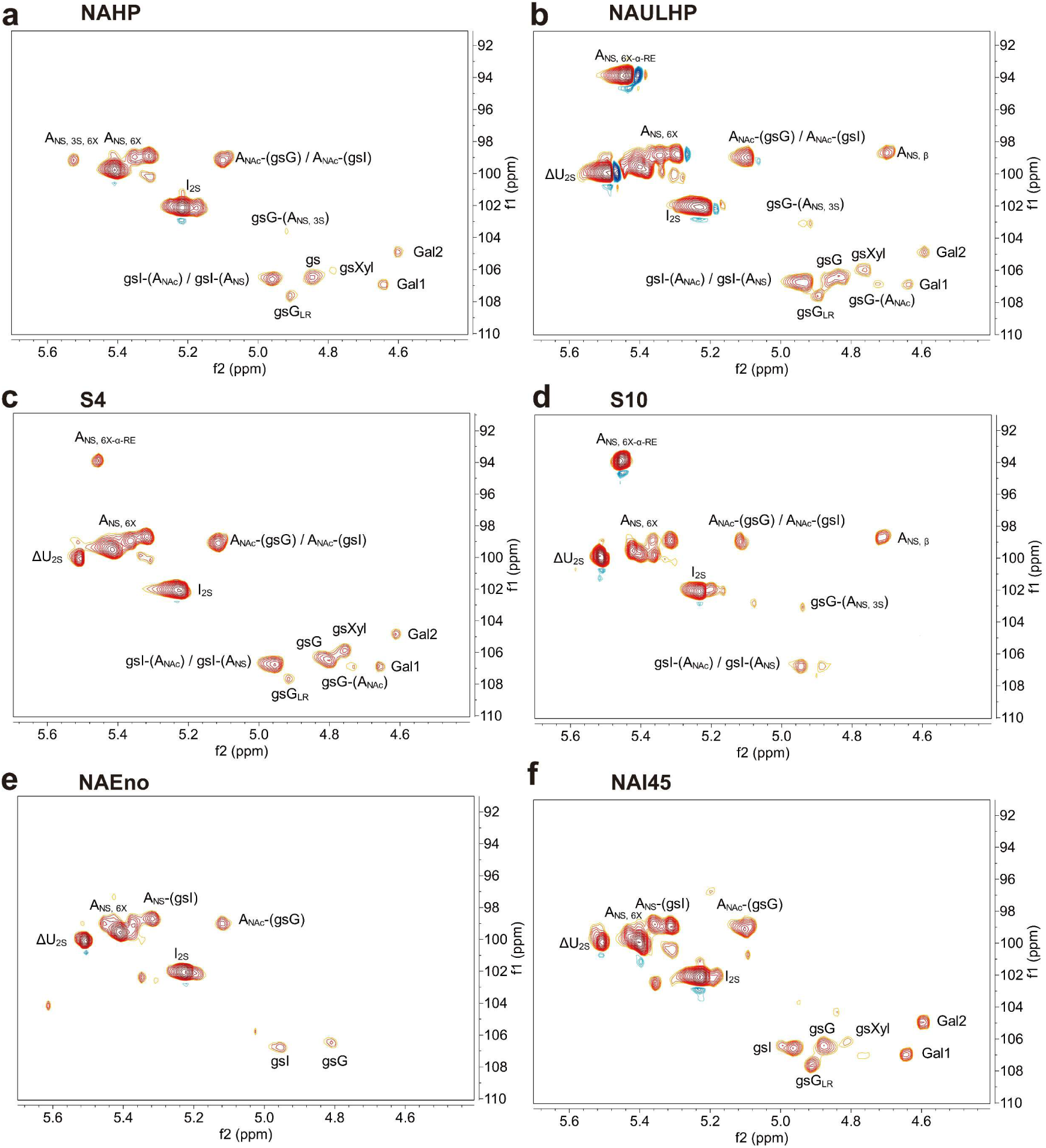
HSQC spectra reveal structural heterogeneity among deanticoagulated heparin derivatives. Representative heteronuclear single-quantum coherence (HSQC) spectra of **a**, NAHP; **b**, NAULHP; **c**, S4; **d**, S10; **e**, NAEno; and **f**, NAI45.

**Supplementary Fig. 8.**
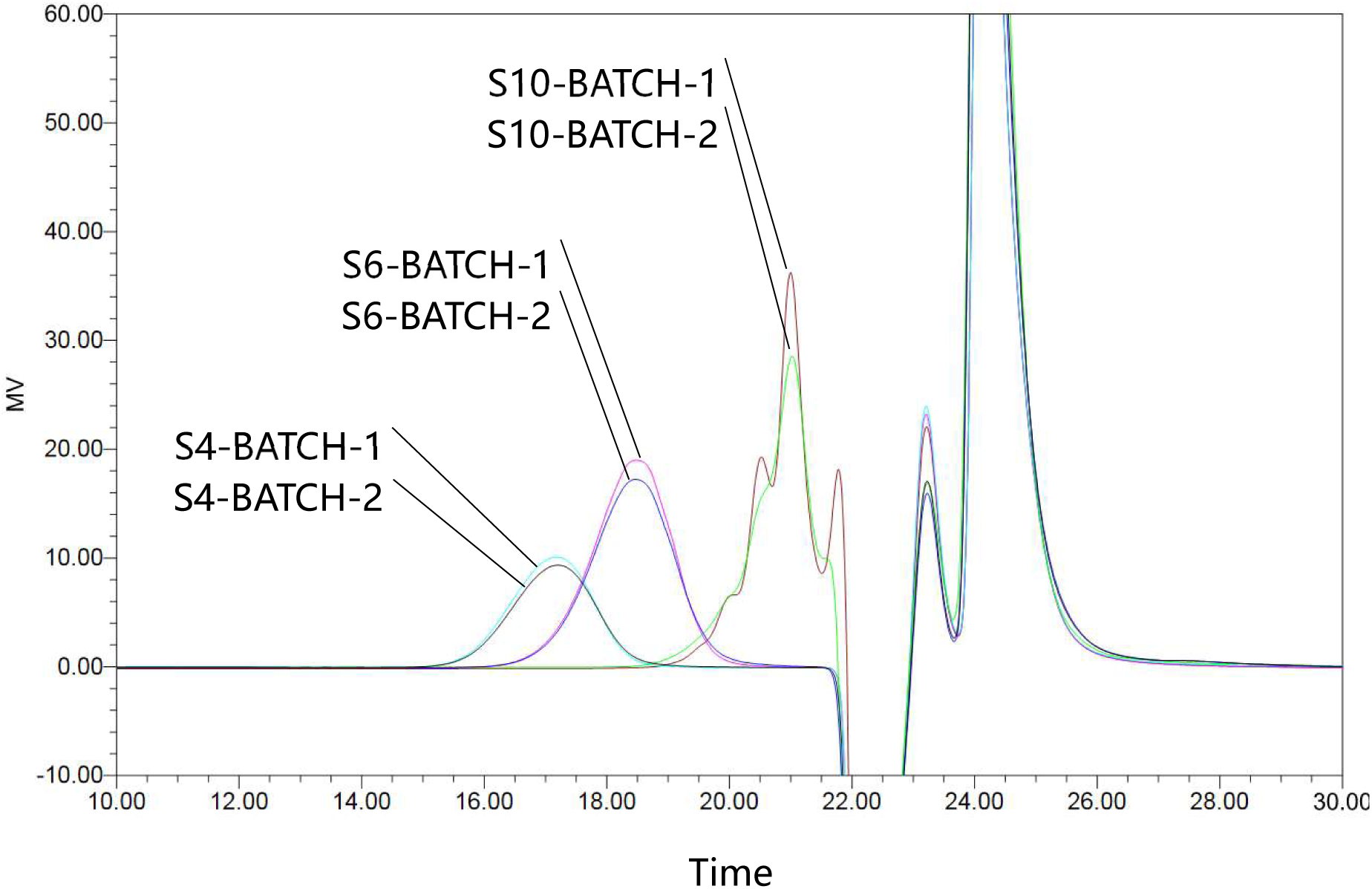
Independent fractionation batches reproducibly isolate S4, S6 and S10 from NALHP. Overlay of chromatographic profiles obtained during two independent preparative separations of S4, S6 and S10 from NALHP.

**Supplementary Fig. 9.**
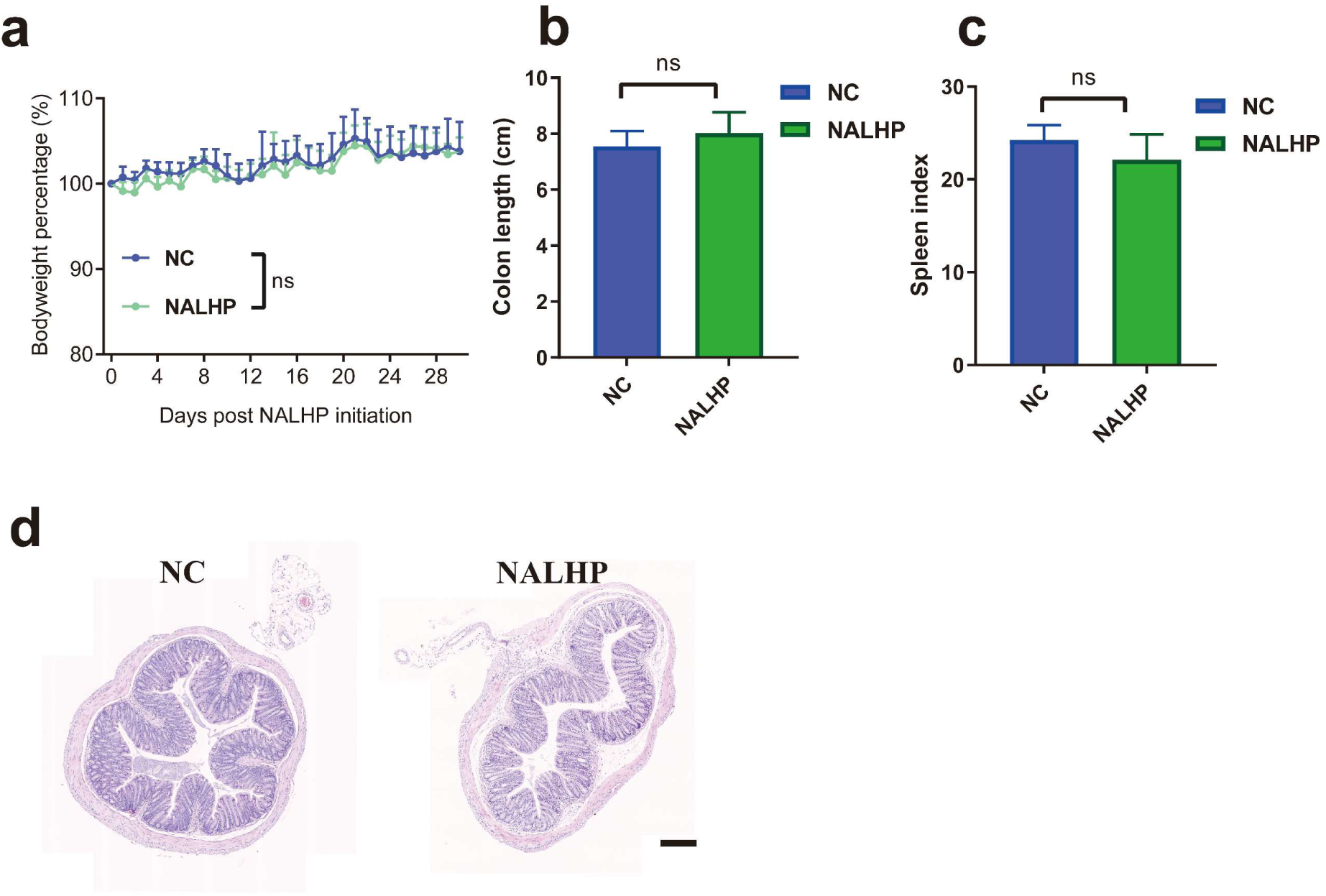
Repeated oral NALHP administration shows no overt intestinal toxicity in healthy mice. **a**, Body-weight changes during 30 days of oral NALHP administration to healthy mice. **b-c**, Colon length (**b**) and spleen index (**c**) at the experimental endpoint. **d**, Representative H&E-stained colon sections from negative-control and NALHP-treated mice. Scale bar, 200 μm. The data are presented as the mean ±SD. * *p* < 0.05, ** *p* < 0.01, *** *p* < 0.001, **** *p* < 0.0001, and ns (not significant) *p* > 0.05, analyzed by ordinary one-way ANOVA multiple comparisons test in (**a**).

**Supplementary Fig. 10.**
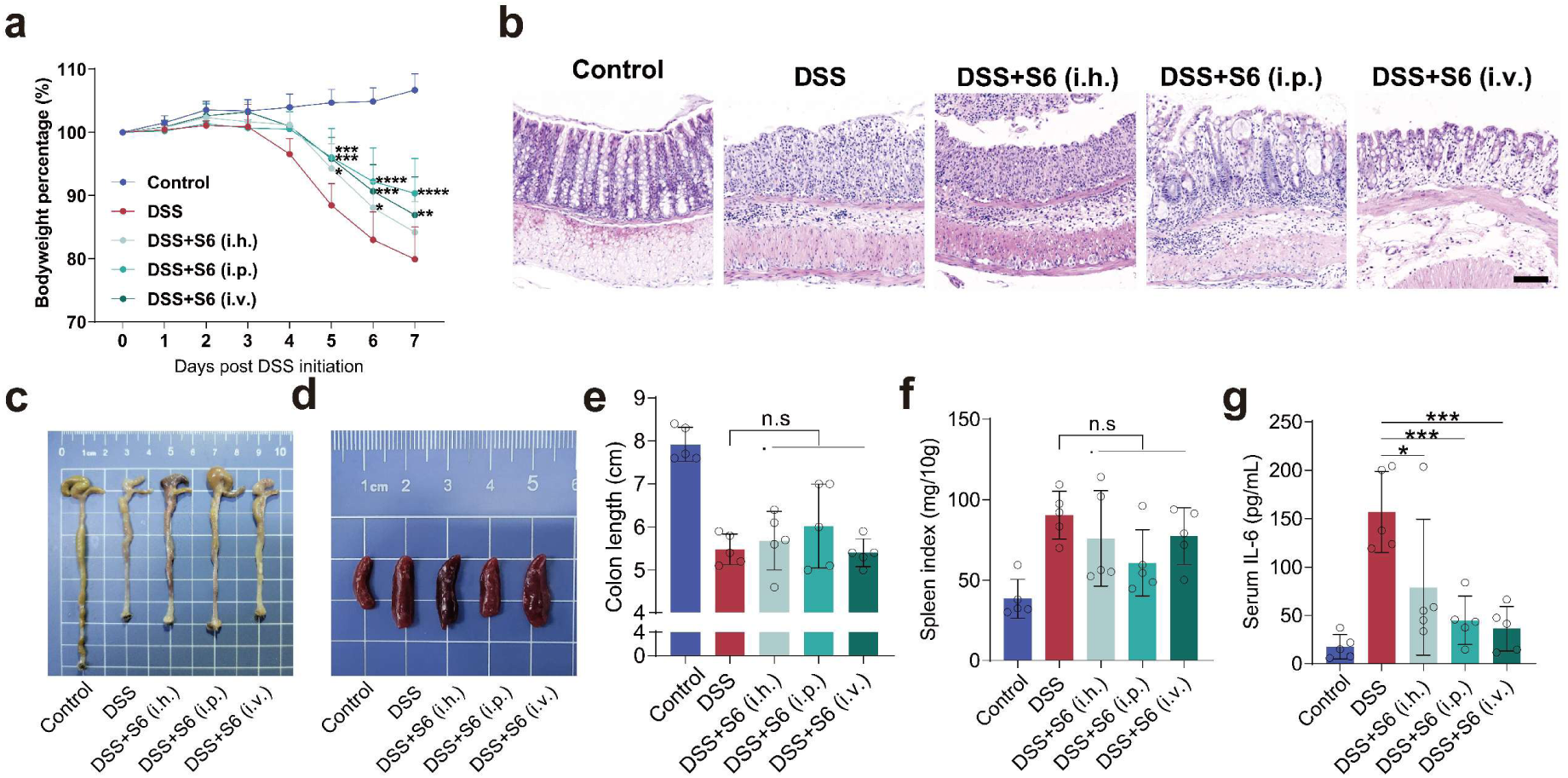
Systemic administration does not improve S6 efficacy relative to the oral route. **a**, Body-weight changes in DSS-treated mice receiving S6 by the indicated administration routes. **b**, Representative H&E-stained colon sections after S6 administration by hypodermic, intraperitoneal or intravenous injection. Scale bar, 100 μm. **c-d**, Representative gross images of colons (**c**) and spleens (**d**). **e-f**, Colon length (**e**) and spleen index (**f**). **g**, Serum IL-6 concentrations. (**a, e-g**) Representative images or quantitative analysis from at least n=5 biologically independent samples are shown. The data are presented as the mean ±SD. * *p* < 0.05, ** *p* < 0.01, *** *p* < 0.001, **** *p* < 0.0001, and ns (not significant) *p* > 0.05, analyzed by ordinary one-way ANOVA multiple comparisons test in (**G**).

**Supplementary Fig. 11.**
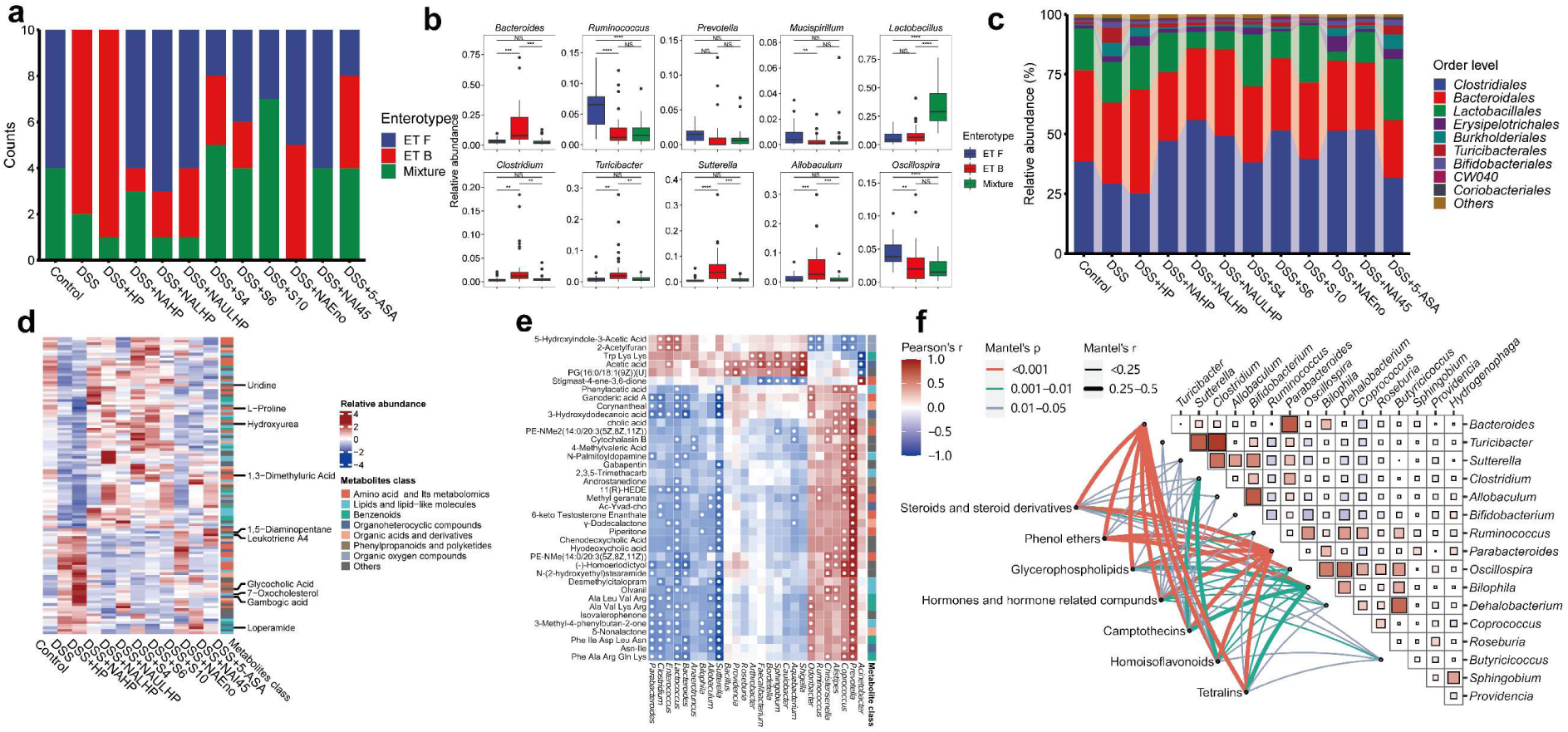
Microbiome and metabolome remodeling does not uniquely track the therapeutic activity of NALHP and S6. **a**, Distribution of intestinal microbial community types following treatment with different heparin derivatives. **b**, Abundance of discriminatory bacterial genera identified by the classification model. **c**, Genus-level microbial community composition across experimental groups. **d**, Heatmap of differentially abundant intestinal metabolites. **e**, Spearman correlation matrix integrating microbial and metabolomic features. **f**, Mantel analysis of associations between microbial taxa and metabolites.

**Supplementary Fig. 12.**
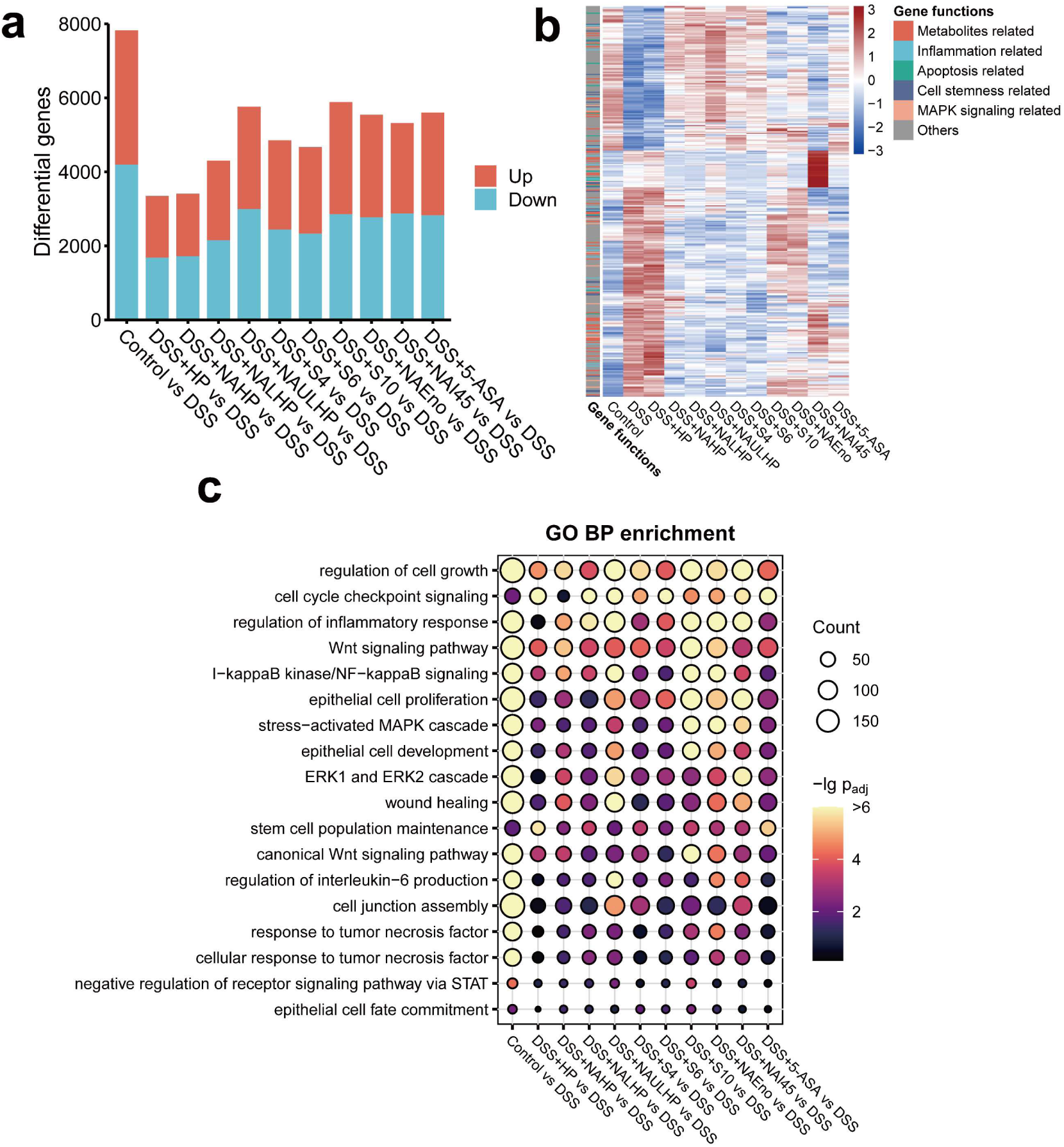
Colonic transcriptomics links therapeutic activity to epithelial proliferation, repair and Wnt-associated programs. **a**, Numbers of differentially expressed genes across pairwise comparisons between DSS-treated mice and the indicated treatment groups. **b**, Heatmap of differentially expressed genes grouped according to major functional categories. **c**, Gene Ontology biological-process enrichment across heparin treatments, highlighting epithelial proliferation, cell-fate regulation, wound healing, cell-junction organization, inflammatory responses and Wnt-associated processes.

**Supplementary Fig. 13.**
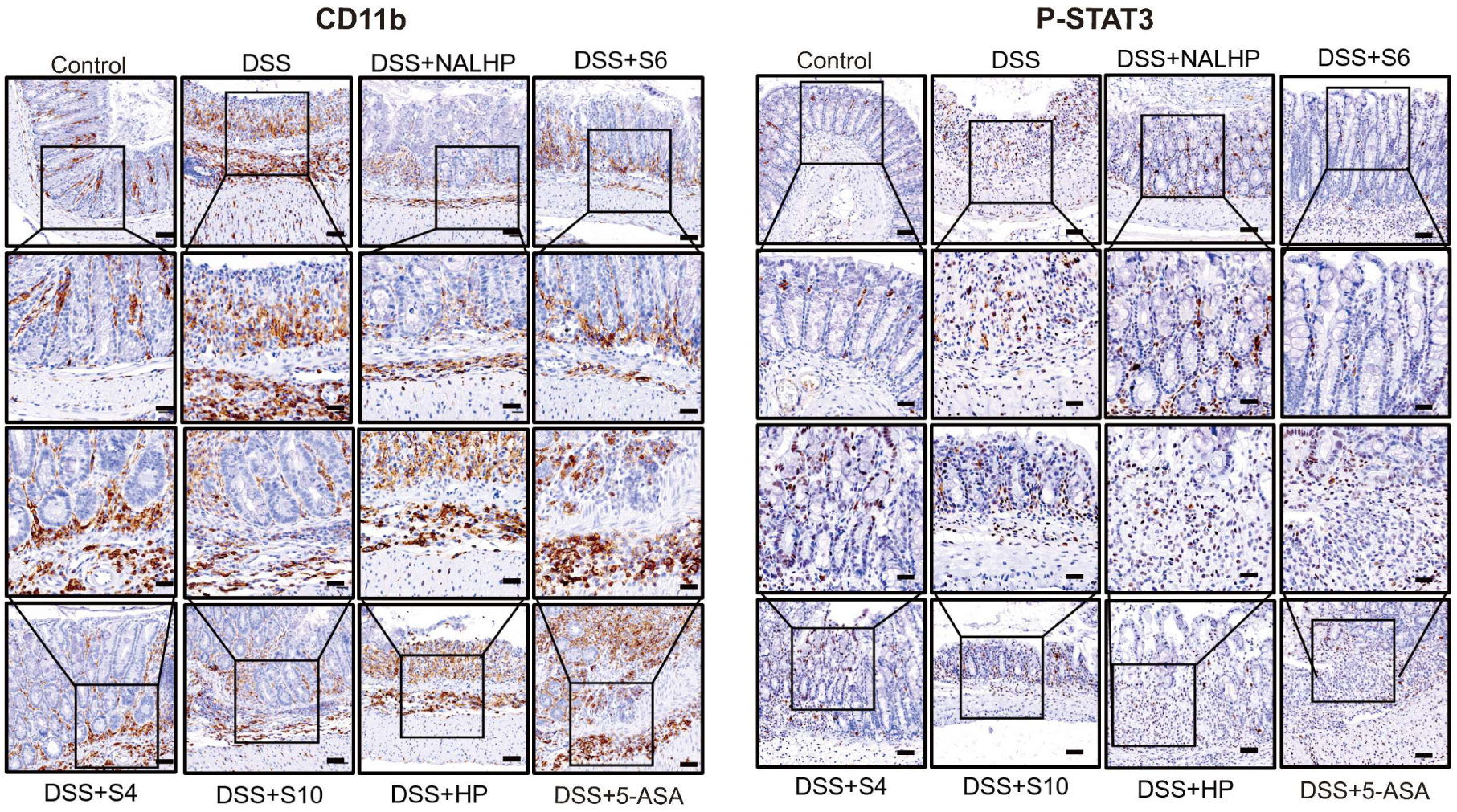
NALHP and S6 reduce local inflammatory infiltration and STAT3 activation in DSS-injured colon. Representative immunohistochemical staining for CD11b and phosphorylated STAT3 (P-STAT3) in colon sections from healthy controls and DSS-treated mice receiving NALHP, S6, S4, S10, HP or 5-ASA. Scale bars, 100 μm. Representative images from n= 5 biologically independent samples are shown.

**Supplementary Fig. 14.**
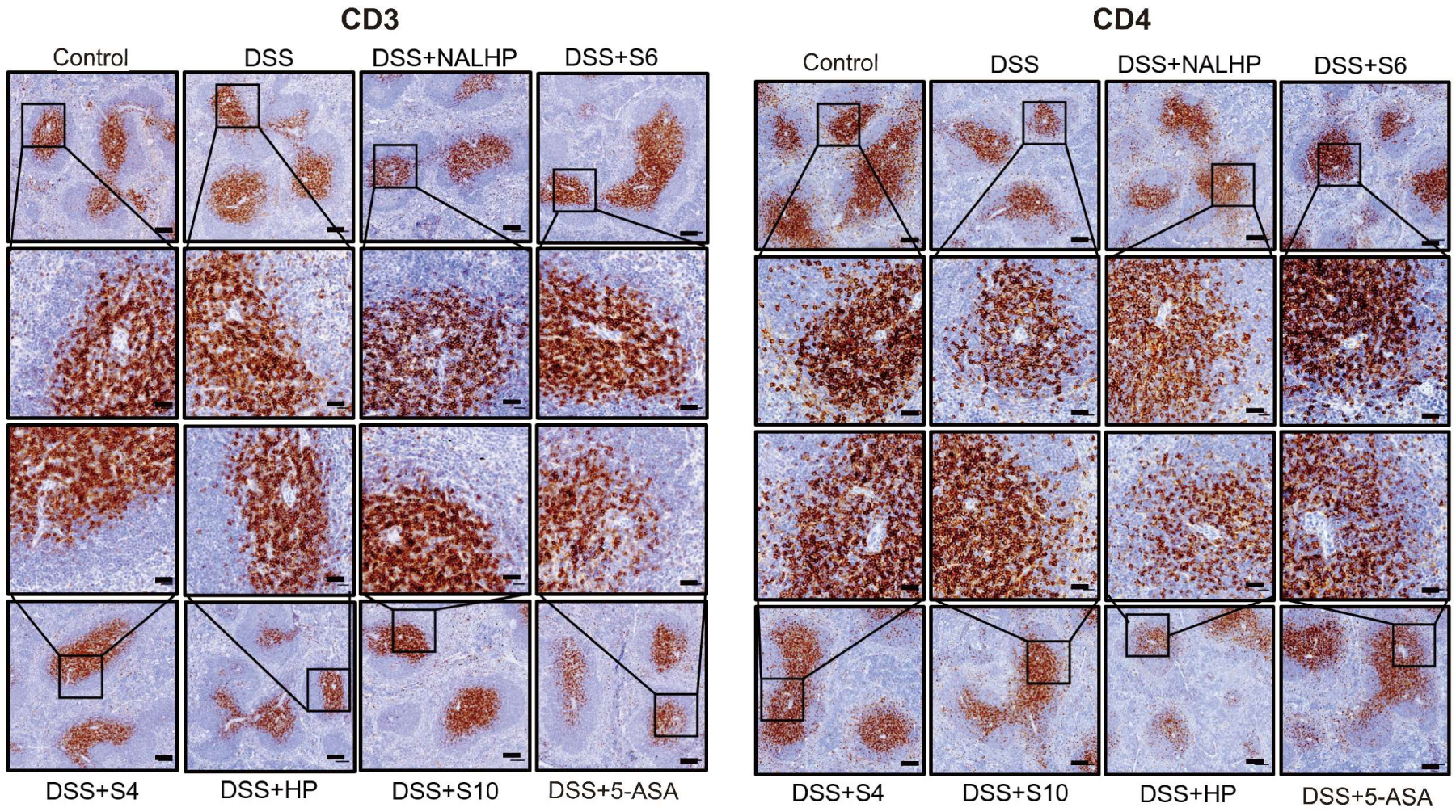
Major splenic T-cell distributions are not substantially remodeled by NALHP/S6 treatment. Representative immunohistochemical staining for CD3 and CD4 in spleens from healthy and DSS-treated mice receiving the indicated treatments. Scale bars, 100 μm. Representative images from n= 5 biologically independent samples are shown.

**Supplementary Fig. 15.**
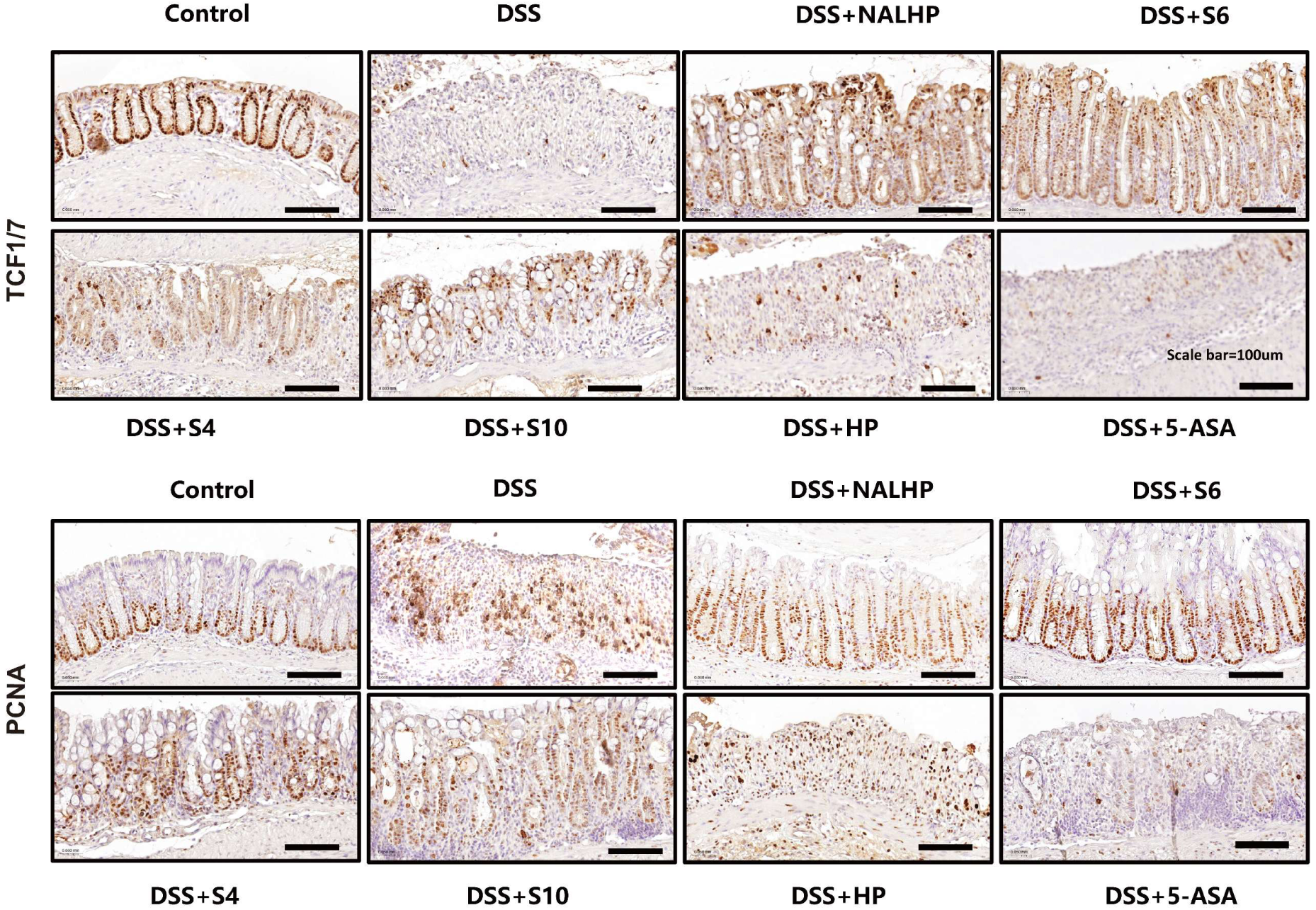
NALHP and S6 normalize crypt proliferative organization and Wnt-associated epithelial signaling in vivo. Representative immunohistochemical staining for TCF1/7 and PCNA in colon sections from healthy controls and DSS-treated mice receiving NALHP, S6, S4, S10, HP or 5-ASA. Scale bars, 100 μm. Representative images from n= 5 biologically independent samples are shown.

**Supplementary Fig. 16.**
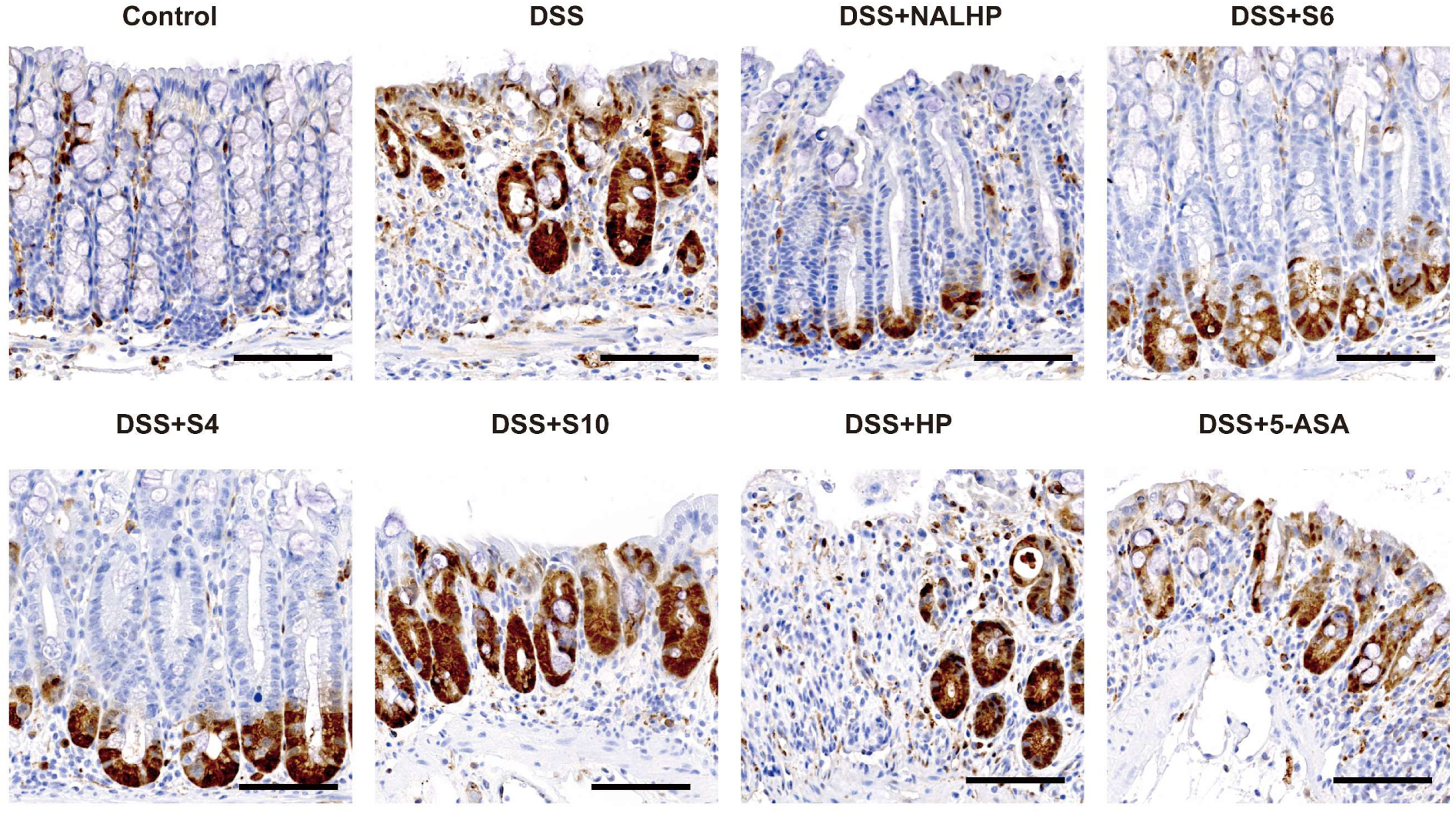
NALHP and S6 remodel ERK-associated crypt signaling during epithelial repair. Representative immunohistochemical staining for phosphorylated ERK (P-ERK) in colon sections from healthy controls and DSS-treated mice receiving the indicated treatments. Scale bars, 100 μm. Representative images from n= 5 biologically independent samples are shown.

**Supplementary Fig. 17.**
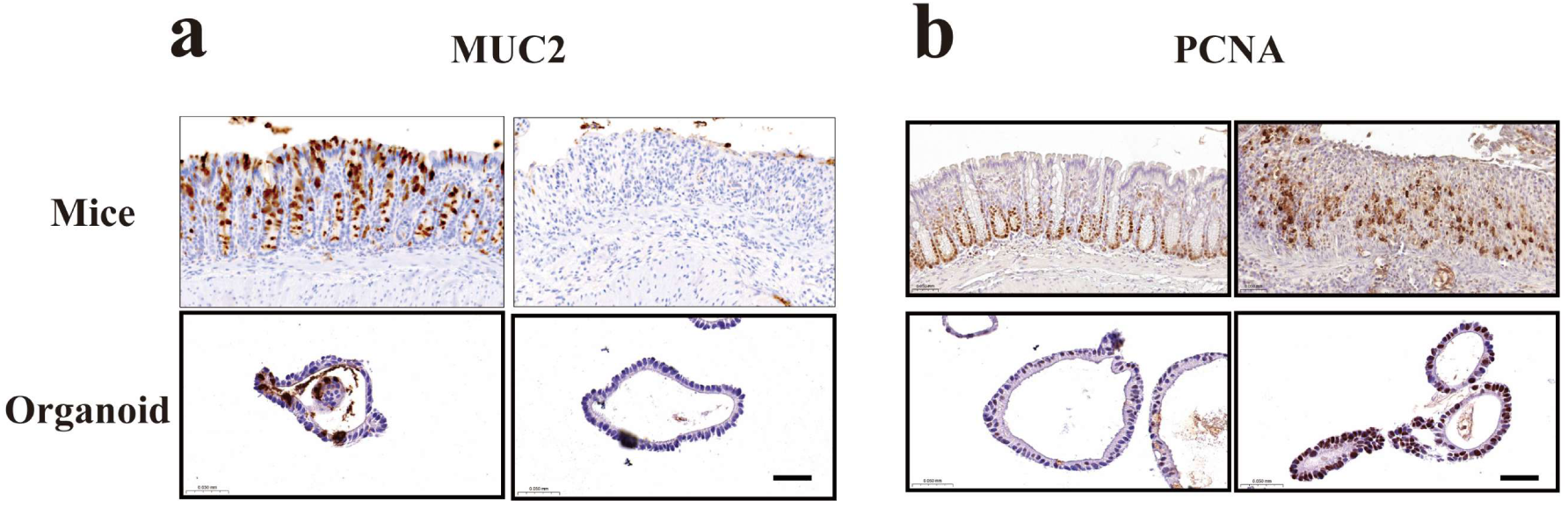
MUC2 and PCNA phenotypes are conserved across UC-associated tissue and organoid models. Representative immunohistochemical staining for MUC2 (**a**) and PCNA (**b**) across human UC-associated samples and experimental mouse colitis, illustrating the reciprocal relationship between mucus-associated differentiation and proliferative epithelial activity. Scale bars, 50 μm. Representative images from n= 3 biologically independent samples are shown.

**Supplementary Fig. 18.**
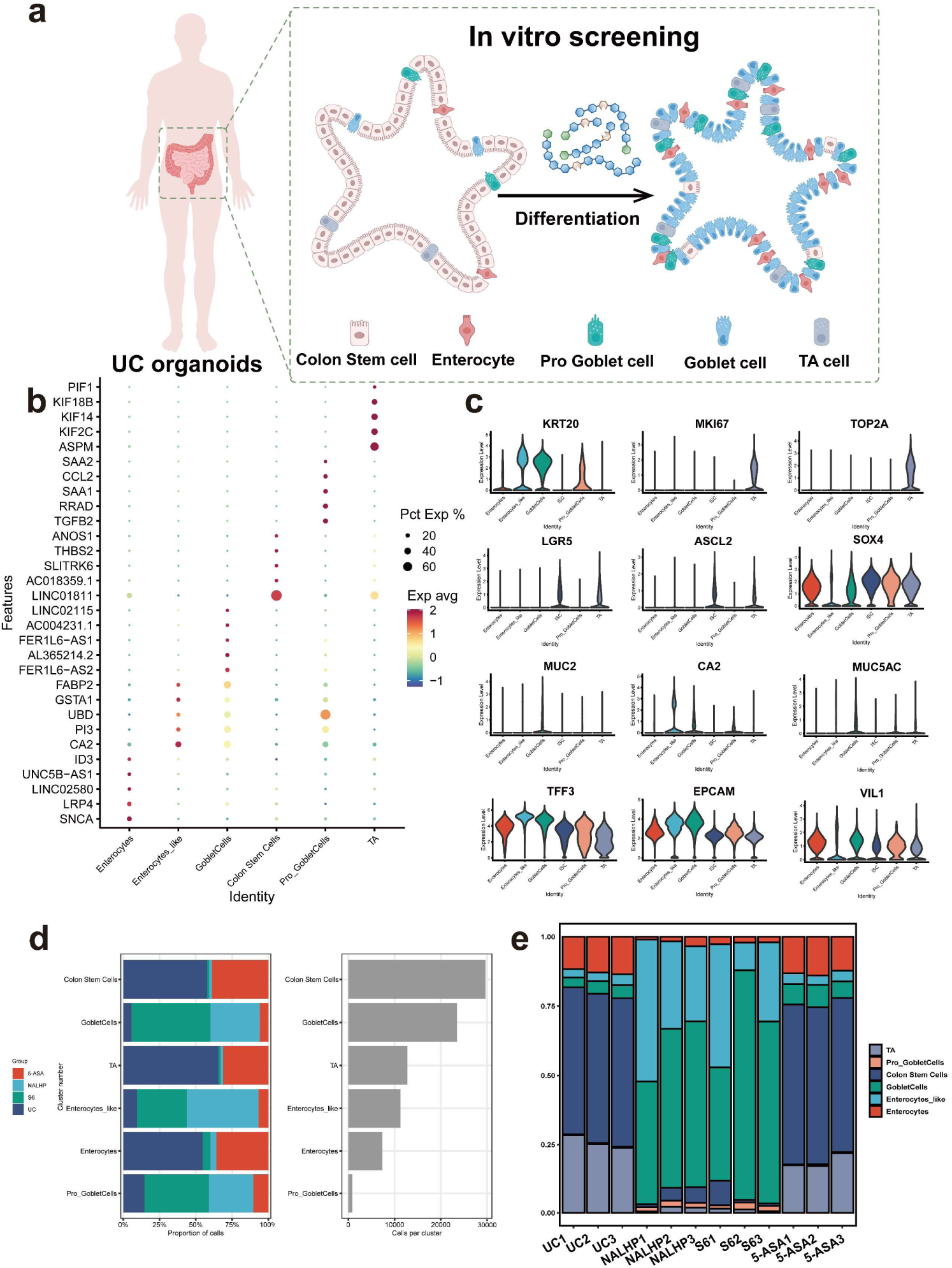
Conventional single-cell annotation confirms broad epithelial remodeling after NALHP/S6 treatment. **a**, Schematic overview of UC patient-derived organoid establishment and epithelial differentiation analysis. **b**, Dot plot of representative markers used for conventional epithelial cell-type annotation. **c**, Expression distributions of selected markers across annotated epithelial populations. **d**, Relative abundance of conventional epithelial cell types across treatment groups. **e**, Sample-level epithelial composition showing reproducibility of treatment-associated remodeling.

**Supplementary Fig. 19.**
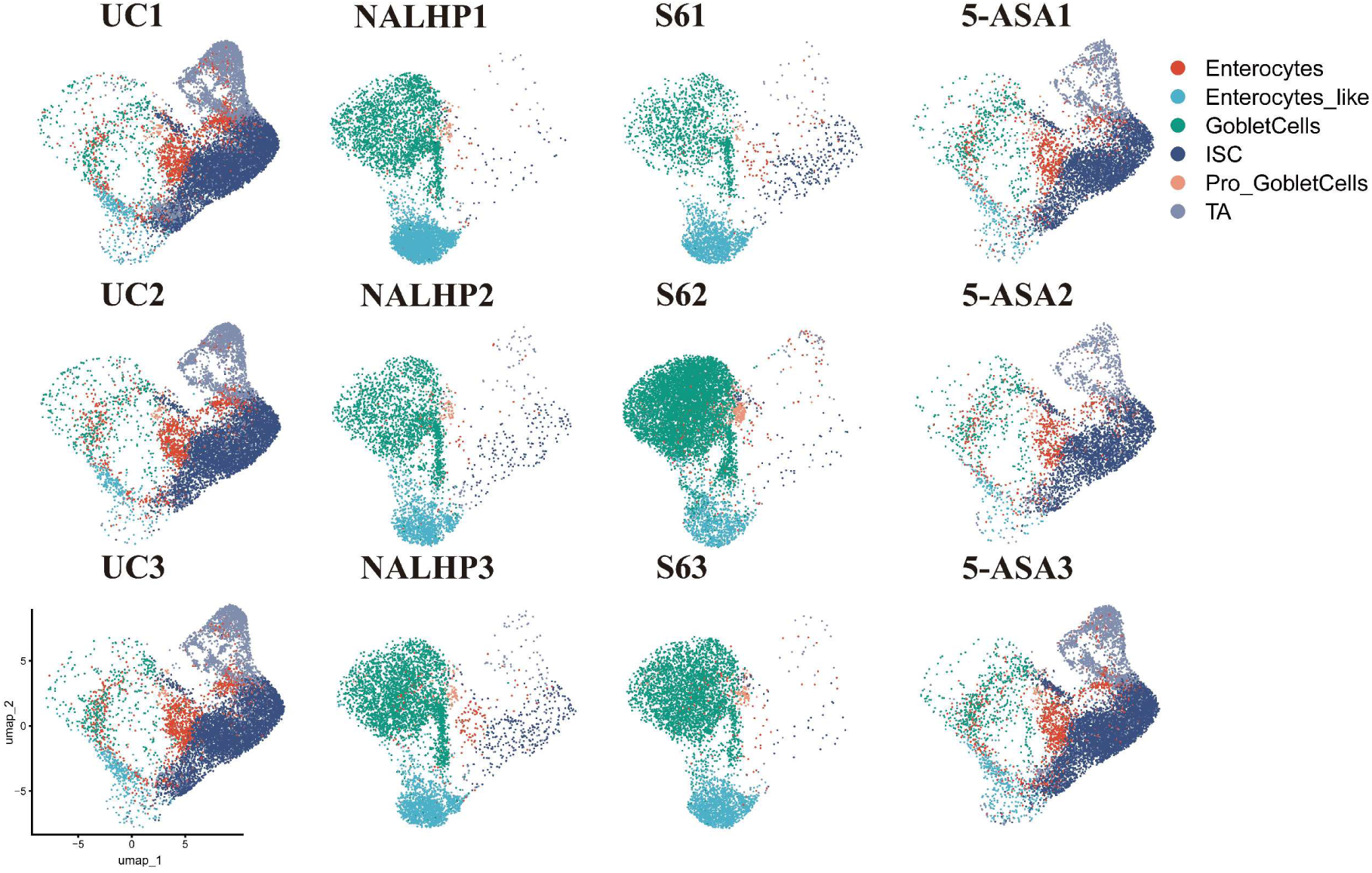
Patient-level single-cell maps confirm reproducible epithelial remodeling across UC organoid lines. UMAP visualizations of epithelial cells from three independently derived UC patient organoid lines (UC1–UC3) under untreated, NALHP-, S6– and 5-ASA-treated conditions. Cells are colored according to conventional epithelial identity.

**Supplementary Fig. 20.**
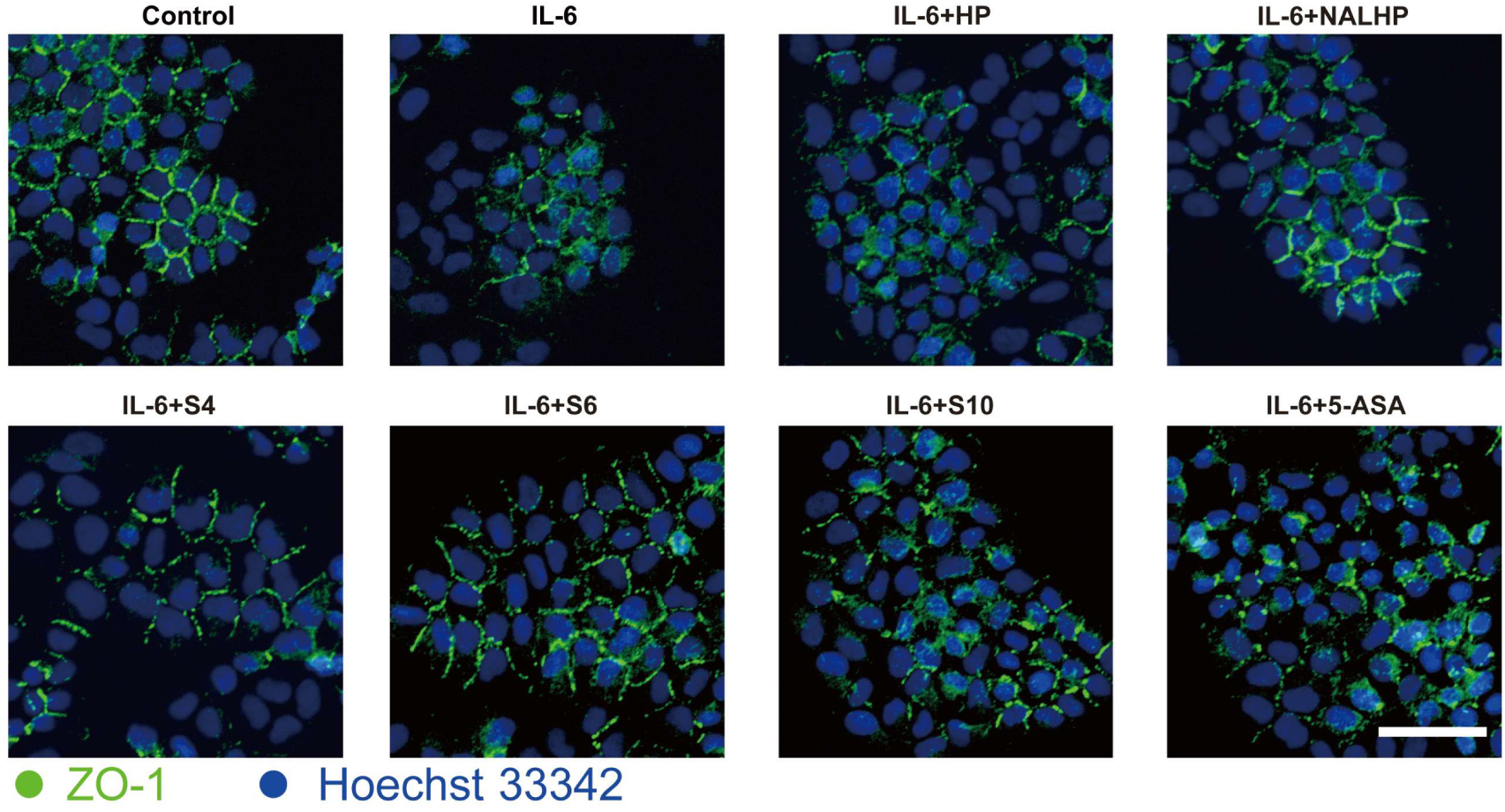
NALHP and selected active heparin fractions restore ZO-1 organization in cytokine-challenged colonic epithelial cells. Representative immunofluorescence images of ZO-1 in NCM460 colonic epithelial cells exposed to IL-6 and subsequently treated with HP, NALHP, S4, S6, S10 or 5-ASA. ZO-1 is shown in green and nuclei are counterstained with Hoechst 33342. Scale bar, 50 μm.

**Supplementary Fig. 21.**
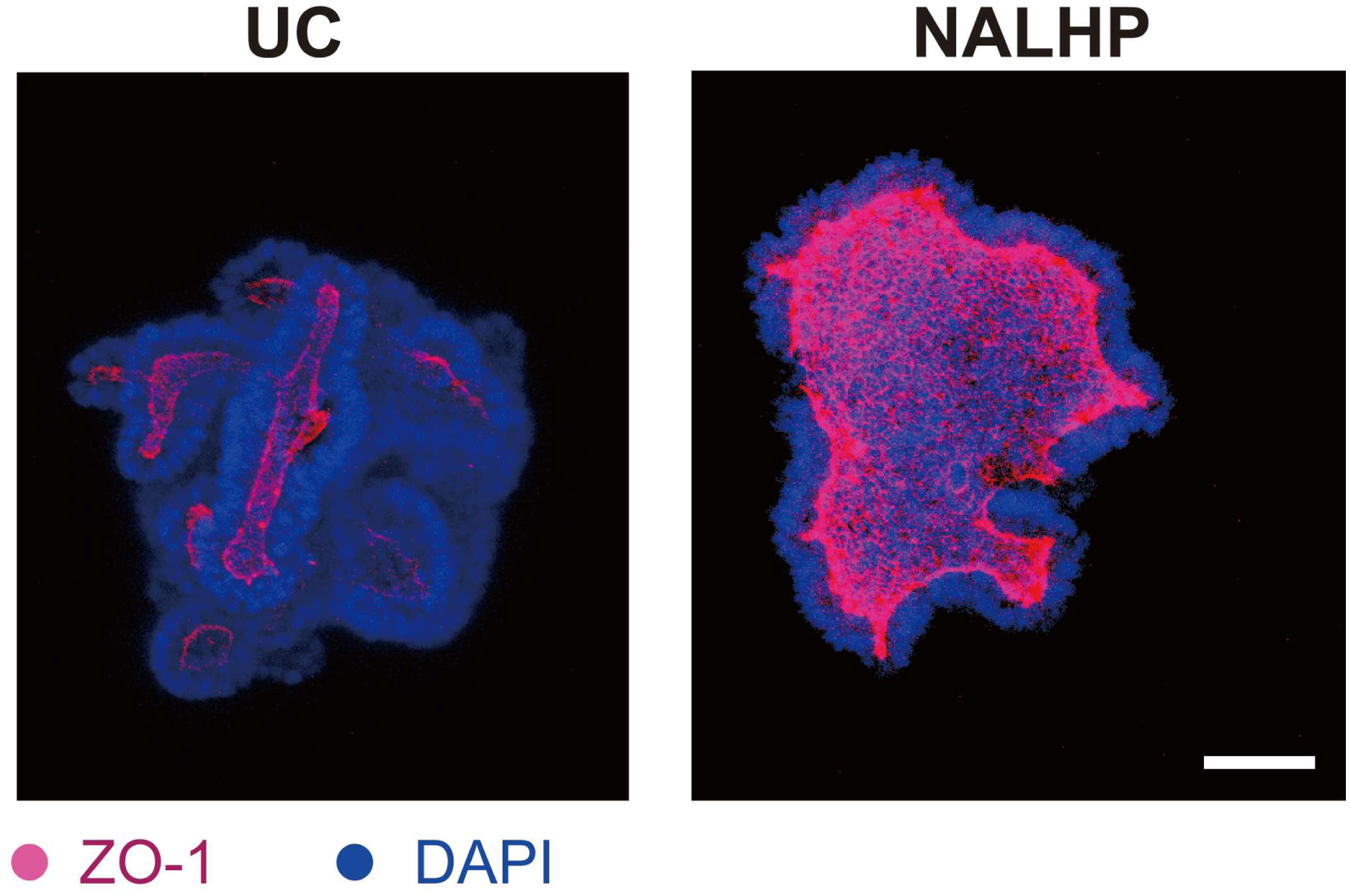
NALHP restores junctional ZO-1 organization in UC patient-derived organoids. Representative immunofluorescence images of ZO-1 in untreated and NALHP-treated UC patient-derived organoids. ZO-1 is shown in magenta and nuclei are counterstained with DAPI. Organoids were treated with NALHP at 10 μg ml⁻¹. Scale bar, 200 μm.

**Supplementary Fig. 22.**
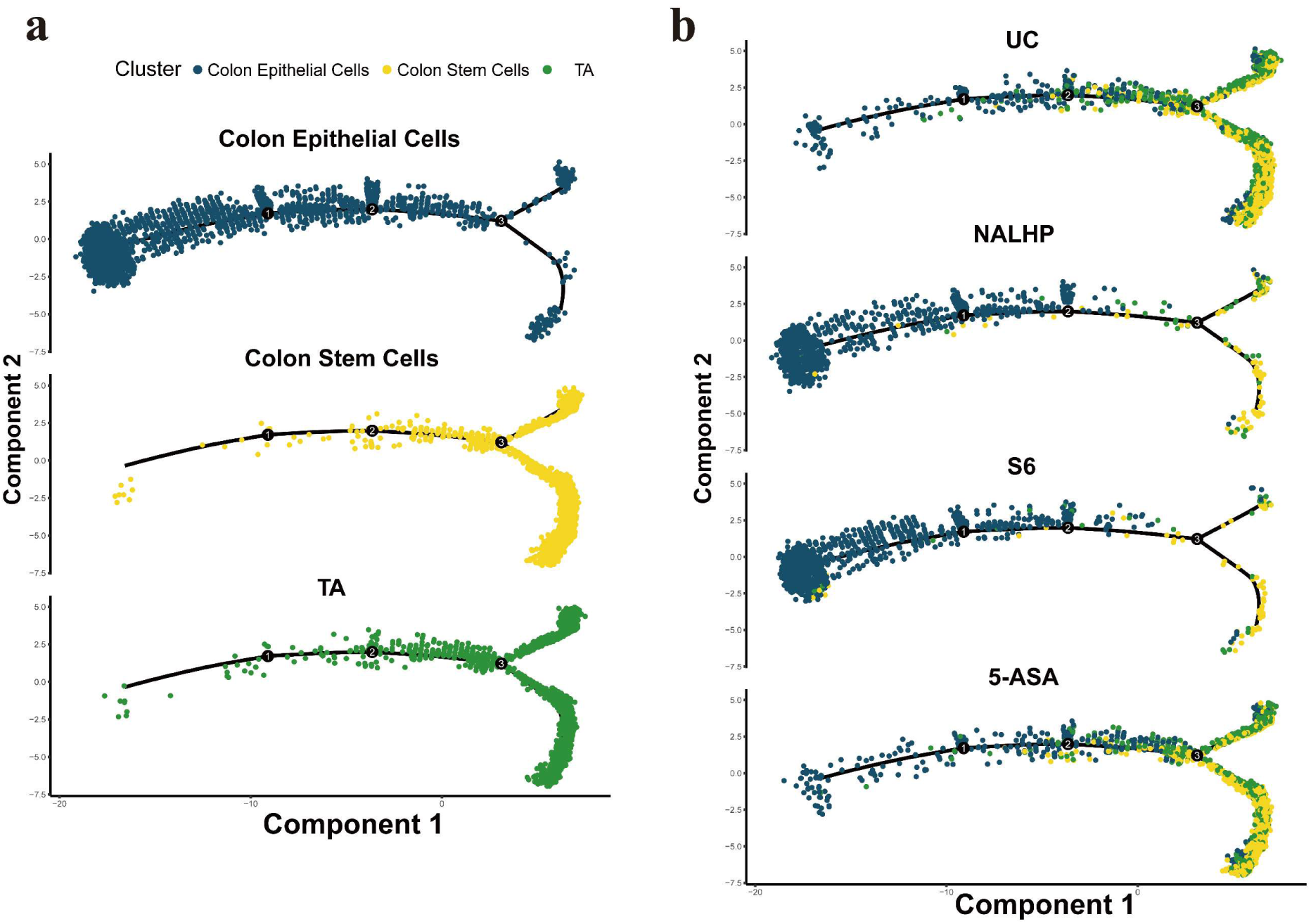
Conventional pseudotime analysis supports treatment-associated progression from stem/proliferative to differentiated epithelial states. **a**, Monocle 2 trajectories visualized according to conventional epithelial annotations, including crypt stem, transit-amplifying and differentiated epithelial populations. **b**, Corresponding trajectories separated by treatment group. The cells were sorted according to pseudotime, and genes were clustered according to their expression patterns (blue: colon epithelial cells; yellow: crypt stem cells; green: TA cells)

**Supplementary Fig. 23.**
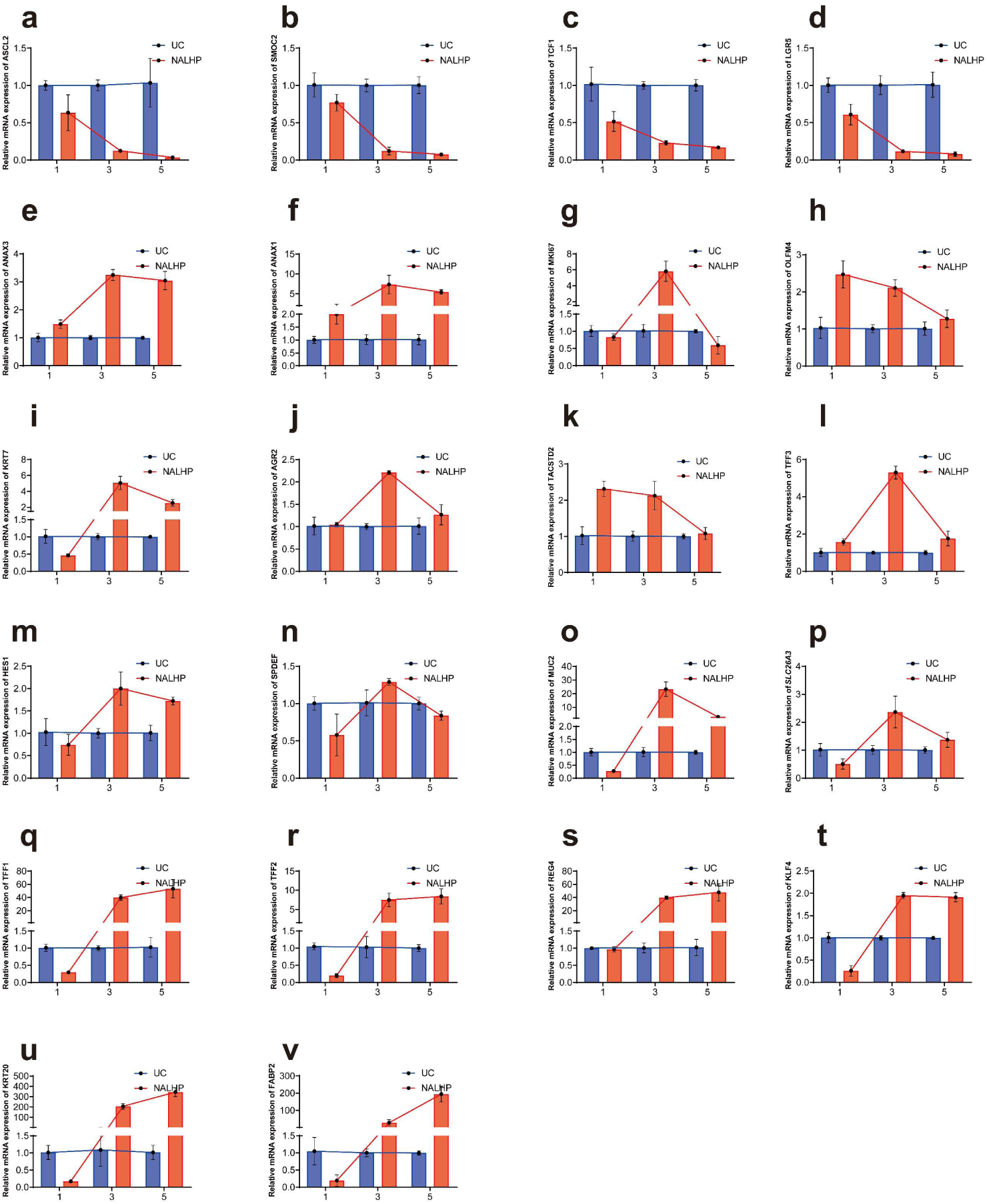
Individual temporal markers support a transient repair-secretory program followed by epithelial maturation. Temporal expression profiles of individual genes associated with classical Wnt-stemness, the repair-secretory transitional state, goblet/mucus-barrier maturation and absorptive maturation in UC organoids treated with NALHP across days 1, 3 and 5. Values are normalized to day-matched untreated UC controls. Stemness-associated genes progressively decline, repair-associated genes exhibit early or intermediate induction, and mature absorptive and mucus-barrier markers become elevated at later time points. These individual-gene profiles complement the state-level temporal analysis in Fig. 4 and support the transient nature of RSTS.

**Supplementary Fig. 24.**
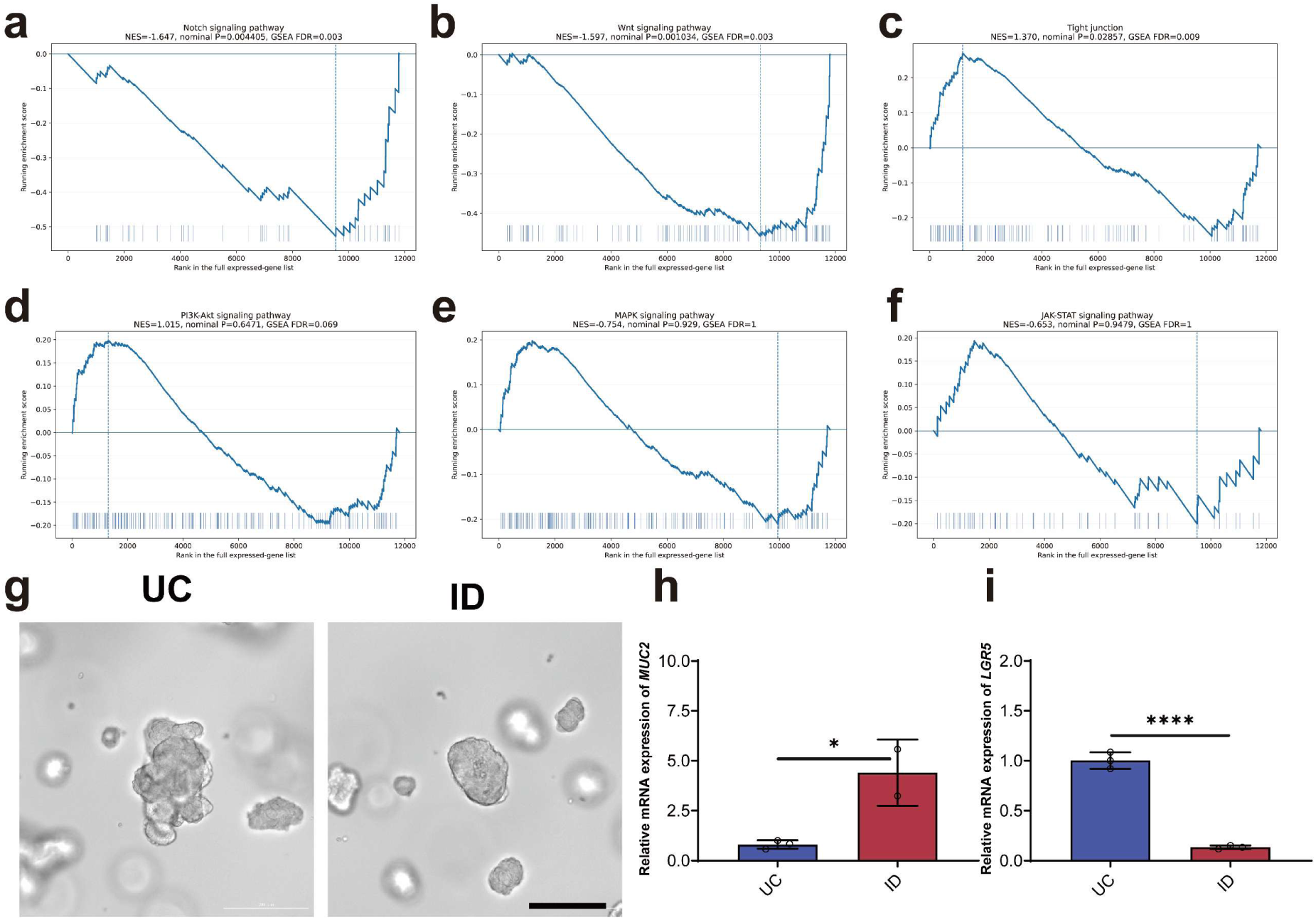
Independent attenuation of Wnt and Notch signaling phenocopies key features of NALHP-induced epithelial maturation. **a–f**, Gene-set enrichment analysis of Notch (**a**), Wnt (**b**), tight-junction (**c**), PI3K–Akt (**d**), MAPK (**e**) and JAK–STAT (**f**) signaling programs in NALHP-treated versus untreated UC organoids. **g**, Representative bright-field images of untreated UC organoids and organoids treated with the Wnt-pathway inhibitor IWP-2 together with the Notch-pathway inhibitor DAPT (ID). Scale bar, 200 μm. **h,i**, Relative mRNA expression of MUC2 (**h**) and LGR5 (**i**) following combined IWP-2/DAPT treatment.

**Supplementary Fig. 25.**
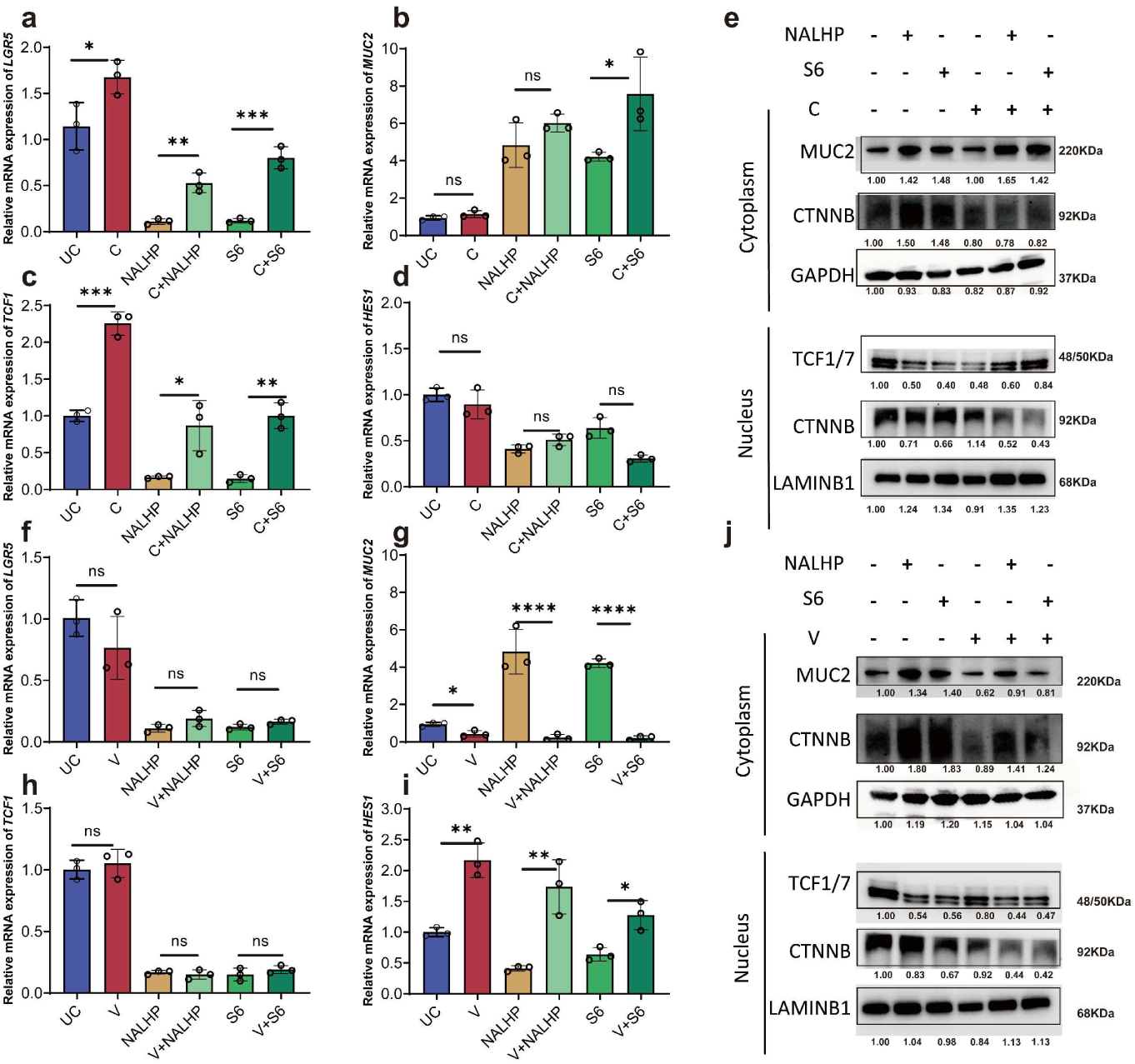
Independent Wnt and Notch reactivation reveals complementary control of NALHP-driven epithelial state progression. **a–d**, Relative mRNA expression of LGR5 (**a**), MUC2 (**b**), TCF1 (**c**) and HES1 (**d**) in UC organoids treated with NALHP or S6 in the absence or presence of the Wnt activator CHIR99021 (C). Wnt reactivation preferentially restored LGR5 and TCF1-associated stemness outputs while exerting a more limited effect on HES1. **e**, Cytoplasmic and nuclear immunoblot analysis of MUC2, CTNNB and TCF1/7 following NALHP/S6 treatment with or without CHIR99021. GAPDH and LAMINB1 served as cytoplasmic and nuclear loading controls, respectively. **f–i**, Relative mRNA expression of LGR5 (**f**), MUC2 (**g**), TCF1 (**h**) and HES1 (**i**) in UC organoids treated with NALHP or S6 in the absence or presence of the Notch activator valproic acid (V). Notch reactivation preferentially restored HES1 and opposed MUC2 induction without comparably restoring the Wnt-associated stemness program. **j**, Cytoplasmic and nuclear immunoblot analysis following NALHP/S6 treatment with or without valproic acid. These results indicate complementary rather than redundant functions of Wnt– and Notch-associated signaling during NALHP-driven epithelial state progression. Data are presented as mean ± SD from *n* = 3 biologically independent experiments. * *p* < 0.05, ** *p* < 0.01, *** *p* < 0.001, **** *p* < 0.0001; ns (not significant) *p* > 0.05.

**Supplementary Fig. 26.**
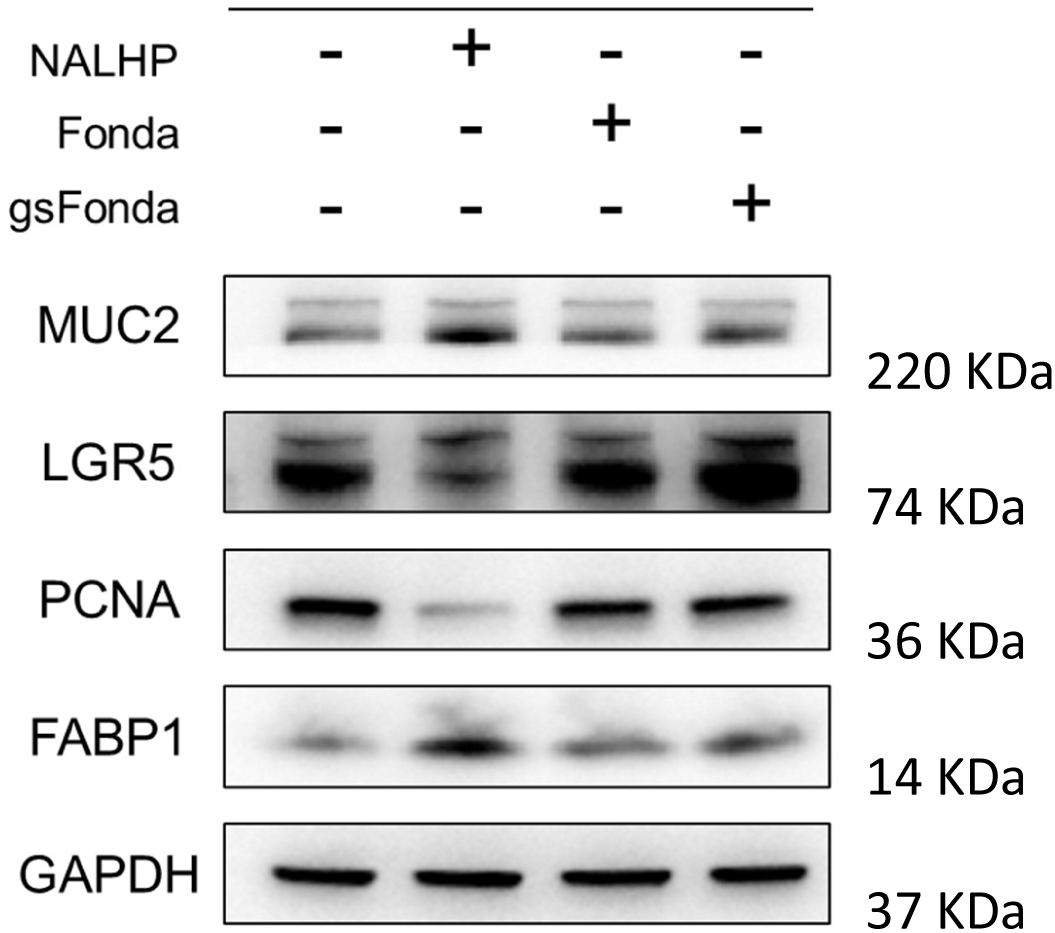
Removal of anticoagulant activity is insufficient to reproduce the epithelial effects of NALHP. Immunoblot analysis of MUC2, LGR5, PCNA and FABP1 in UC patient-derived organoids treated with NALHP, fondaparinux (Fonda) or non-anticoagulated fondaparinux (gs-Fonda). GAPDH served as the loading control.

**Supplementary Table. 1.** Molecular weight and size distribution of NALHP and its fine fragments.

| Name | Mass<br>(mg) | Content<br>(%, w/w) | *M<sub>w</sub>* (Da) | *P* | *M<sub>w</sub>* (Da) distribution (%) |  |  |  |
| --- | --- | --- | --- | --- | --- | --- | --- | --- |
|  |  |  |  |  | >8k | 5k~8k | 3k~5k | <3k |
| NALHP | 650 | 100 | 6224 | 2.33 | 28.04 | 27.87 | 19.11 | 24.95 |
| NAULHP | - | - | 2974 | 3.22 | 5.21 | 14.96 | 20.05 | 59.77 |
| S1 | 37.7 | 5.8 | 12121 | 1.036 | 99 | 1 |  |  |
| S2 | 44.2 | 6.8 | 9557 | 1.037 | 80 | 20 |  |  |
| S3 | 42.9 | 6.6 | 7686 | 1.038 | 38 | 60 | 2 |  |
| S4 | 37.7 | 5.8 | 6254 | 1.041 | 8 | 78 | 13 |  |
| S5 | 89.7 | 13.8 | 5094 | 1.055 | 1 | 50 | 46 | 2 |
| S6 | 99.5 | 15.3 | 4115 | 1.076 |  | 19 | 68 | 13 |
| S7 | 71.5 | 11.0 | 3227 | 1.117 |  | 5 | 50 | 45 |
| S8 | 78.65 | 12.1 | 2426 | 1.172 |  | 1 | 23 | 76 |
| S9 | 62.4 | 9.6 | 1736 | 1.188 |  |  | 7 | 93 |
| S10 | 48.1 | 7.4 | 1282 | 1.153 |  |  | 1 | 99 |
| S11 | 37.7 | 5.8 | 812 | 1.073 |  |  |  | 100 |

**Supplementary Table. 2.** Characteristic infrared peaks of heparin-like polysaccharides.

| Wavenumber<br>(cm<sup>-1</sup>) | Source of the characteristic<br>peak | Type of peak |
| --- | --- | --- |
| 800 | IdoA<u>2S</u> | C-O-S stretching and vibration |
| 820 | GlcNS<u>6S</u> | C-O-S stretching and vibration |
| 890 | Glc<u>NS</u> | C-O-S stretching and vibration,<br>C-O-C vibration coupling |
| 940 | GlcNS<u>6S</u> | C-O-S stretching and vibration,<br>C-O-C glycosidic bond |
| 1230 | Glc<u>NS</u> | S=O asymmetric stretching vibration |
| 1378 | GlcN<u>Ac</u> | -CH<sub>3</sub> vibration |
| 1560 | GlcN<u>Ac</u> | Amide bond |

**Supplementary Table. 3.** Batch numbers of the raw materials and products for the 4 production batches of NALHP.

| Batch Number | #1 | #2 | #3 | #4 |
| --- | --- | --- | --- | --- |
| HP | NHS210204 | NHS200606 | NHS200803 | NHS180302 |
| NAHP | BATCH-211019 | BATCH-210401 | BATCH-210319 | BATCH-190304 |
| NALHP | BATCH-211023 | BATCH-210408 | BATCH-210402 | BATCH-190504 |

**Supplementary Table. 4.** Preparation process parameter standards for the 4 production batches of NALHP.

| Process Indicator | #1 | #2 | #3 | #4 | Process<br>standard | Qualified<br>or<br>unqualified |
| --- | --- | --- | --- | --- | --- | --- |
| anti-Xa (IU/mg) | 19.4 | 18.1 | 18.5 | 19.0 | <20 | Qualified |
| $A_{232}$ | 19.0 | 12.2 | 28.1 | 23.4 | <30 | Qualified |
| $\Delta A_{232}$ | 41.0 | 39.7 | 38.8 | 39.6 | 39.5±2.0 | Qualified |
| $A_{294}$ | 0.167 | 0.134 | 0.189 | 0.245 | <0.350 | Qualified |
| $A_{420}$ | 0.018 | 0.019 | 0.015 | 0.022 | <0.025 | Qualified |
| NALHP $\Delta M_w$ (Da) | 102 | 136 | 150 | 242 | 300~300 | Qualified |

**Supplementary Table. 5.**
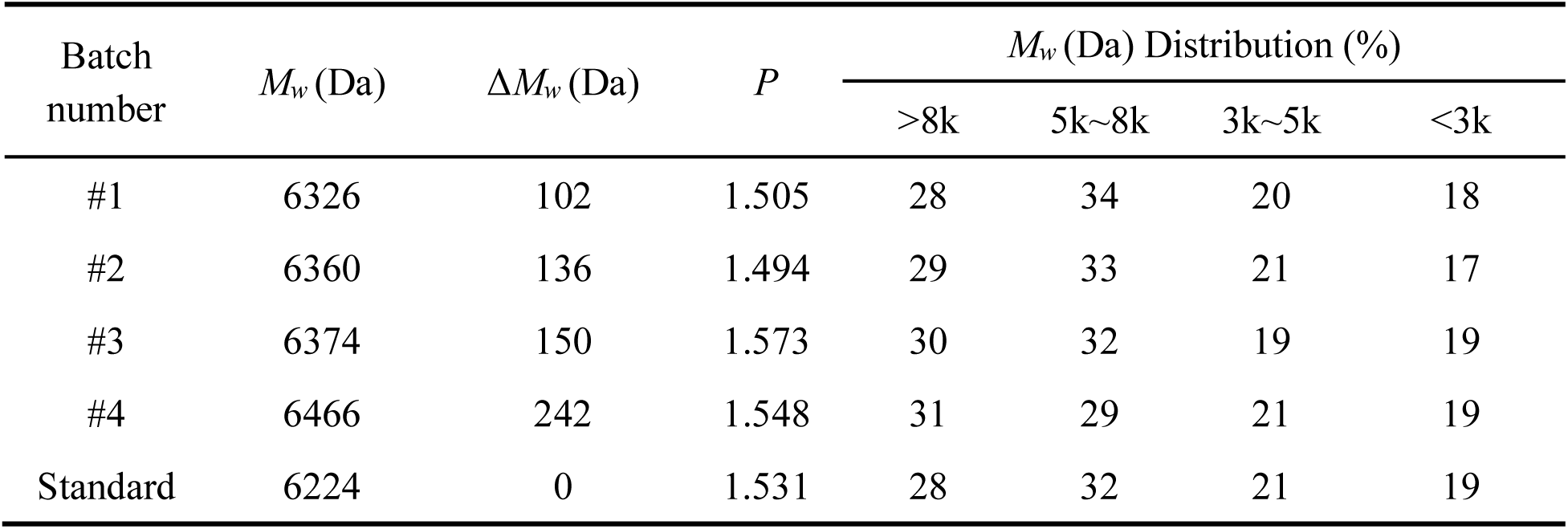
Molecular weight and size distribution of the 4 production batches of NALHP.

**Supplementary Table. 6.** ^13^C NMR chemical shifts of HP and different deanticoagulated HPs and their assignments.

| Peak position | Chemical shift of the sample (ppm) |  |  |  |  |  |  | Chemical shift assignment |
| --- | --- | --- | --- | --- | --- | --- | --- | --- |
|  | HP | NAHP | NALHP | NAULHP | S4 | S6 | S10 |  |
| a | 177.7 | 176.7 | 177.7 | 177.7 | 177.7 | 177.7 | 177.7 | U, -COOH;<br>GlcNAc6S/GlcNAc, CH<sub>3</sub>CO- |
| b | - | - | 172.1 | 172.1 | 172.1 | 172.1 | 172.1 | ΔU/ΔU2S, -COOH |
| c | - | - | 147.6 | 147.6 | 147.6 | 147.6 | 147.6 | ΔU2S, **C5** |
| d | - | - | 108.9 | 108.9 | 108.9 | 108.9 | 108.9 | ΔU2S, **C4** |
| e | - | 106.7 | 106.7 | 106.7 | 106.7 | 106.7 | 106.7 | gsIdoA/gsGlcA, **C1** |
| f | 102.3 | 102.3 | 102.3 | 102.3 | 102.3 | 102.3 | 102.3 | IdoA2S, **C1** |
| g | 99.9 | 99.9 | 99.9 | 99.9 | 99.9 | 99.9 | 99.9 | ΔU2S/GlcNS6S/GlcNS/GlcNAc6S/GlcNAc, **C1** |
| h | - | 99.1 | 99.1 | 99.1 | 99.1 | 99.1 | 99.1 | **GlcNAc-**<br>(gsIdoA)/**GlcNAc-**<br>(gsGlcA), **C1** |
| i | - | - | 93.9 | 93.9 | 93.9 | 93.9 | 93.9 | GlcNS-α-RE |
| j | 24.8 | 24.8 | 24.8 | 24.8 | 24.8 | 24.8 | 24.8 | -CH<sub>3</sub>CO- |

**Supplementary Table. 7.** ^1^H NMR chemical shifts of HP and different deanticoagulated HPs and their assignments.

| Peak position | Chemical shift of the sample (ppm) |  |  |  |  |  |  | Chemical shift assignment |
| --- | --- | --- | --- | --- | --- | --- | --- | --- |
|  | HP | NAHP | NALHP | NAULHP | S4 | S6 | S10 |  |
| a | - | - | 5.99 | 5.99 | 5.99 | 5.99 | 5.99 | ΔU, **H4** |
| b | 5.58 | - | - | - | - | - | - | **GlcNS-(GlcA), H1** |
| c | 5.53 | 5.53 | 5.53 | 5.53 | 5.53 | 5.53 | 5.53 | GlcNS3S6S, **H1** |
| d | - | - | 5.52 | 5.52 | 5.52 | 5.52 | 5.52 | ΔU, **H1** |
| e | 5.42 | 5.42 | 5.42 | 5.42 | 5.42 | 5.42 | 5.42 | GlcNS6S/GlcNAc6X/GlcN<br>**S6X-(gsIdoA)/GlcNS6X-**<br>(gsGlcA), **H1** |
| f | 5.35 | - | - | - | - | - | - | **GlcNS6X-(IdoA), H1** |
| g | 5.22 | 5.22 | 5.22 | 5.22 | 5.22 | 5.22 | 5.22 | IdoA2S, **H1** |
| h | - | 5.12 | 5.12 | 5.12 | 5.12 | 5.12 | 5.12 | **GlcNAc-(gsIdoA)/GlcNAc-**<br>(gsGlcA), **H1** |
| i | 5.01 | - | - | - | - | - | - | **IdoA-(GlcNS), H1** |
| j | 4.83 | 4.92 | 4.83 | 4.83 | 4.83 | 4.83 | 4.83 | IdoA2S, **H5** |
| k | - | 4.97 | 4.97 | 4.97 | 4.97 | 4.97 | 4.97 | gsIdoA, **H1** |
| l | - | 4.85 | 4.85 | 4.85 | 4.85 | 4.85 | 4.85 | gsGlcA, **H1** |
| m | - | 4.73 | 4.73 | 4.73 | 4.73 | 4.73 | 4.73 | gsXyl, **H1** |
| n | 4.64 | 4.64 | 4.64 | 4.64 | 4.64 | 4.64 | 4.64 | Gal, **H1** |
| o | 4.61 | - | - | - | - | - | - | GlcA, **H1** |
| p | - | - | 4.63 | 4.63 | 4.63 | 4.63 | 4.63 | ΔU, **H2** |
| q | 4.34 | 4.34 | 4.34 | 4.34 | 4.34 | 4.34 | 4.34 | IdoA2S, **H2** |
| r | 3.40 | - | - | - | - | - | - | GlcA, **H2** |
| s | 3.28 | 3.28 | 3.28 | 3.28 | 3.28 | 3.28 | 3.28 | GlcNS6X, **H2** |
| t | 2.05 | 2.05 | 2.05 | 2.05 | 2.05 | 2.05 | 2.05 | -CH<sub>3</sub>CO |

**Supplementary Table. 8.** Patient information.

| Group | Age | Gender | Sampling site | Mayo score |
| --- | --- | --- | --- | --- |
| UC1 | 35 | Male | Colon | 5 |
| UC2 | 30 | Male | Colon | 7 |
| UC3 | 29 | Female | Colon | 6 |

**Supplementary Table. 9.**
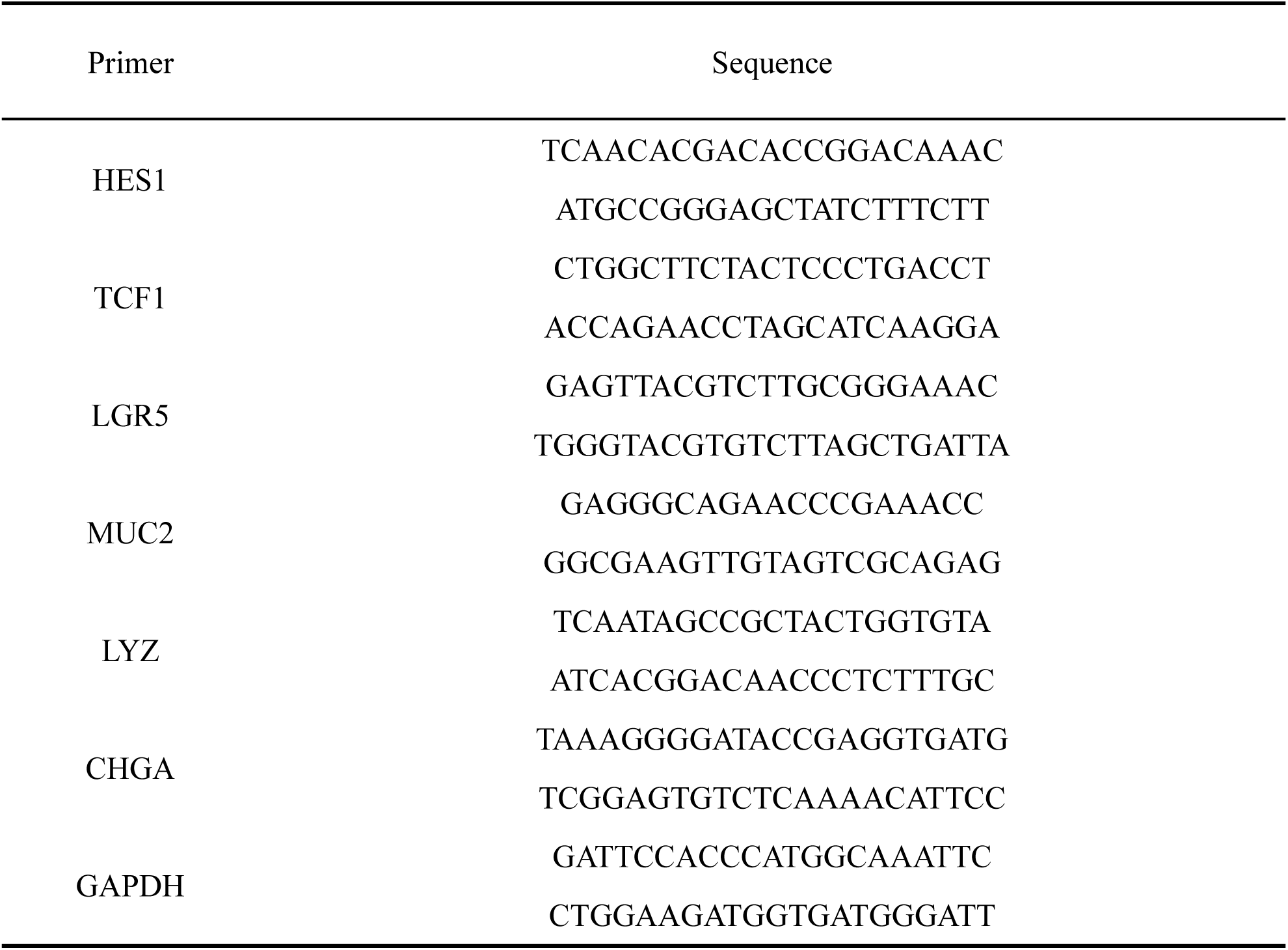
qPCR primer sequences.

**Supplementary Table. 10.** Antibodies and reagents.

| Antibody | Company | CatLog | Application | Dilution rate |
| --- | --- | --- | --- | --- |
| MUC2 | Abcam | ab272692 | WB/IHC | 1:2000/1:200 |
| PCNA | CST | 2586 | WB/IHC | 1:10000/1:2000 |
| TCF1/7 | CST | C63D9 | WB/IHC | 1:2000/1:200 |
| CTNNB | CST | 8480 | WB | 1:2000 |
| GAPDH | CST | 5174 | WB | 1:2000 |
| LAMINB1 | CST | 13435 | WB | 1:1000 |
| Anti-CD4 | CST | 25229 | IHC | 1:100 |
| Anti-phospho ERK1/2<br>(Thr202/Tyr204) | CST | 4370 | IHC | 1:400 |
| Anti-phospho STAT3<br>(Tyr705) | CST | 9145 | IHC | 1:200 |
| Anti-WGA Lectin | GeneTex | GTX01502 | IF | 1:250 |
| Anti-F4/80 | CST | 70076 | IHC | 1:250 |
| Anti-mouse IgG | CST | 7076 | WB | 1:10000 |
| Anti-rabbit IgG | CST | 7074 | WB | 1:10000 |

Supplementary materials are available for this paper.

